# NMNAT promotes glioma growth through regulating post-translational modifications of p53 to inhibit apoptosis

**DOI:** 10.1101/2021.05.12.443736

**Authors:** Jiaqi Liu, Xianzun Tao, Yi Zhu, Chong Li, Kai Ruan, Zoraida Diaz-Perez, Hongbo Wang, R. Grace Zhai

## Abstract

Gliomas are highly malignant brain tumors with poor prognosis and short survival. NAD^+^ has been shown to impact multiple processes that are dysregulated in cancer; however, anti-cancer therapies targeting NAD^+^ synthesis have been unsuccessful due to insufficient mechanistic understanding. Here we adapted a *Drosophila* glial neoplasia model and discovered the genetic requirement for NAD^+^ synthase nicotinamide mononucleotide adenylyltransferase (NMNAT) in glioma progression *in vivo* and in human glioma cells. Overexpressing enzymatically active NMNAT significantly promotes glial neoplasia growth and reduces animal viability. Mechanistic analysis suggests that NMNAT interferes with DNA damage-p53-caspase-3 apoptosis signaling pathway by enhancing NAD^+^-dependent posttranslational modifications (PTMs) poly(ADP-ribosyl)ation (PARylation) and deacetylation of p53. Interestingly, NMNAT forms a complex with p53 and PTM enzyme PARP1 to facilitate PARylation. As PARylation and deacetylation reduce p53 pro-apoptotic activity, our results demonstrate that NMNAT promotes glioma progression through regulating p53 post-translational modifications. Our findings reveal a novel tumorigenic mechanism involving protein complex formation of p53 with NAD^+^ synthetic enzyme NMNAT and NAD^+^-dependent PTM enzymes that regulates glioma growth.

## INTRODUCTION

Glioma is the most common intrinsic tumor of the central nervous system deriving from the neoplastic glial cells or neuroglia (Goodenberger & Jenkins, 2012). Based on pathological criteria, gliomas are classified from WHO grade I to IV, among which the high-grade gliomas generally have a much poorer prognosis (Wesseling & Capper, 2018). Several major cellular signaling pathways associated with glioma have been well studied, including RTK/Ras/PI3K, p53, and RB signaling pathways (Cancer Genome Atlas Research, 2008). In addition, metabolism factors, such as IDH1/2, were found to play important roles in glioma (Yan et al., 2009). IDH1 is an enzyme of tricarboxylic acid (TCA) cycle in glucose metabolism and the main producer of NADPH (Molenaar, Radivoyevitch, Maciejewski, van Noorden, & Bleeker, 2014). However, drugs targeting these pathways showed a limited clinical response, indicating a critical need for the mechanistic understanding of the metabolic requirement for glioma tumorigenesis.

Nicotinamide adenine dinucleotide (NAD^+^) is an essential signaling cofactor that regulates cancer metabolism through its co-enzymatic function for many bioenergetic pathways including glycolysis, TCA cycle, and oxidative phosphorylation (Hanahan & Weinberg, 2000). Multiple processes associated with NAD^+^ signaling are dysregulated in cancer, including DNA repair, cell proliferation, differentiation, and apoptosis (Chiarugi, Dolle, Felici, & Ziegler, 2012). Inherited polymorphisms and epigenetic repression of DNA damage repair genes are significantly correlated with the risk of gliomas, indicating that abnormal DNA damage repair plays important roles in glioma formation and progression (Chen, Shao, Chen, Kwan, & Chen, 2010; L. Qi et al., 2017). One of the key initiation events of DNA damage response is poly (ADP-ribose) polymerase (PARP)-mediated poly(ADP-ribosyl)ation (PARylation), the main process that consumes nuclear NAD^+^ (Ame, Spenlehauer, & de Murcia, 2004). Moreover, NAD^+^-dependent SIRTs-mediated deacetylation regulates many oncogenes and tumor suppressor genes in cancer cells (Brooks & Gu, 2009). Consistently, a high level of NAD^+^ is observed in gliomas (Reddy et al., 2008; Tso et al., 2006), and 90% of gliomas are susceptible to NAD^+^ deletion (Tateishi et al., 2015). Therefore, it is critical for rapidly proliferating glioma cells to replenish the NAD^+^ pool for survival.

In the past years, targeting NAD^+^ metabolism has been considered for cancer therapy, and most efforts have been focused on nicotinamide phosphoribosyltransferase (NAMPT), the rate-limiting enzyme of the NAD^+^ salvage pathway, whose expression is increased in multiple types of cancer (Garten et al., 2015; Lucena-Cacace, Otero-Albiol, Jimenez-Garcia, Munoz-Galvan, & Carnero, 2018; Ohanna et al., 2018; Pylaeva et al., 2019). Disappointingly, several clinical trials of NAMPT inhibitors have failed due to low efficacy and high toxicities (Sampath, Zabka, Misner, O’Brien, & Dragovich, 2015), which demands the urgent consideration of an alternative target in the NAD^+^ metabolic pathway. Nicotinamide mononucleotide adenylyltransferase (NMNAT), the last enzyme in the NAD^+^ salvage synthetic pathway, has recently emerged as a potential candidate (Chiarugi et al., 2012). NMNAT has three isoforms in mammals with distinct subcellular localizations: NMNAT1, in the nucleus; NMNAT2, in the cytosol; and NMNAT3, in the mitochondria (Berger, Lau, Dahlmann, & Ziegler, 2005). Dysregulations of both NMNAT1 and NMNAT2 have been implicated in cancer. For example, NMNAT1 is considered a poor prognostic marker for renal cancer (Uhlen et al., 2015; Uhlen et al., 2017). Decreased NMNAT1 expression leads to epigenetic silencing of tumor suppressor genes (Henderson, Miranda, & Emerson, 2017). Inhibition of NMNAT1 delays DNA repair and increases rRNA transcription (Song et al., 2013). In colorectal cancer, NMNAT2 upregulation correlates with the cancer invasive depth and TNM stage (Cui et al., 2016; J. Qi et al., 2018). In non-small cell lung cancer (NSCLC), the NMNAT2 enzymatic activity is upregulated by SIRT3-mediated deacetylation process or p53 signaling (H. Q. Li et al., 2013; Pan et al., 2014). Moreover, depletion of NMNAT2 inhibits cell growth indirectly through reducing glucose availability in neuroblastoma cells (Ryu et al., 2018). These observations indicate the regulatory link between compartmentalized NAD^+^ synthesis and cellular metabolism and rapid cancer cell growth, and further underscore the potential of NMNAT as a viable alternative target in NAD^+^ synthetic pathway, given their aberrant regulation and critical role in cancer metabolism.

In this report, to address the knowledge gap of the role of NMNAT in glioma, we adapted an *in vivo* glial neoplasia in *Drosophila* (Read, Cavenee, Furnari, & Thomas, 2009) and discovered the genetic requirement for NMNAT in glioma growth. Combined with a human glioma cell culture model, we characterized the mechanism of NMNAT in glioma progression. Our results identified the upregulation of enzymatically active NMNAT as an essential metabolic regulator for promoting gliomagenesis and revealed the tumorigenic mechanism of NMNAT-sustained PARylation and deacetylation of p53 in apoptosis suppression.

## RESULTS

### NMNAT is upregulated in oncogenic *Ras^v12^* induced glial neoplasia

The Ras/Raf/ERK signaling cascade is one of the most conserved pathways both in *Drosophila* and human, which is part of the MAP kinase signaling network mainly respond to stress activators (Morrison, 2012). *RAS* mutations in human cancer have long been recognized, with the most common in *KRAS* (85%), and much less in *NRAS* (12%) and *HRAS* (3%) (Simanshu, Nissley, & McCormick, 2017). Upregulated RAS and mutant *RAS* have been detected in glioma which indicates Ras is required for glioma growth (Arvanitis et al., 1991; Guha, Feldkamp, Lau, Boss, & Pawson, 1997; Knobbe, Reifenberger, & Reifenberger, 2004; Rajasekhar et al., 2003). Activation of Ras has been used to model human glioma in *Drosophila* (Read, 2011; Read et al., 2009).

*Ras* oncogene at 85D (*Ras85D*) is the *Drosophila* orthologue of human *RAS*. The constitutively active *Ras85D* mutation (G12V), *Ras^v12^*, has been suggested to be analogous to human oncogenic *RAS* mutation and used to induce tumor (Barbacid, 1987; Wu, Pastor-Pareja, & Xu, 2010). We established a *Drosophila* glial neoplasia model by overexpressing *Ras^v12^* in glial cells, driven by the pan-glial driver repo-GAL4 (Read et al., 2009). Green fluorescent protein (GFP) was co-expressed as a reporter to mark the Ras expressing cells. Under normal conditions, the *Drosophila* central nervous system (CNS) is wrapped by perineurial, subperineurial, and ensheathing glia (Freeman, 2015). Powered with high-resolution quantitative brain morphology analysis (Brazill, Zhu, Li, & Zhai, 2018), we analyzed glial neoplasia tissue using three criteria, i) tissue double-positive for GFP and endogenous Repo expression; ii) tissue mass consists of multiple layers of glia of at least 400 cells, and iii) tissue mass volume greater than 12.4 x 10^3^ μm^3^ (Figure 1-figure supplement 1). When *Ras^v12^* was expressed in glia, numerous glial neoplasia tissues marked by GFP and Repo in the brain and ventral nerve cord (VNC) were detected as early as 100 hours after egg laying (AEL) and the volumes of glial neoplasia increased with age (Fig. 1A, B and G). The brain tumors caused early lethality in pupal stage and greatly reduced survival rate (Fig. 1H). Notably, compared with the normal brain (Fig. 1C and E), we found significantly increased endogenous NMNAT in glial cells at both 100 and 150 hours AEL. The increases of NMANT were most prominent in the nuclear region (Fig. 1D and F), suggesting a possible role for nuclear NMNAT in *Ras^v12^*-induced glial neoplasia formation and progression in *Drosophila*.

**Figure 1 with 1 supplement.**
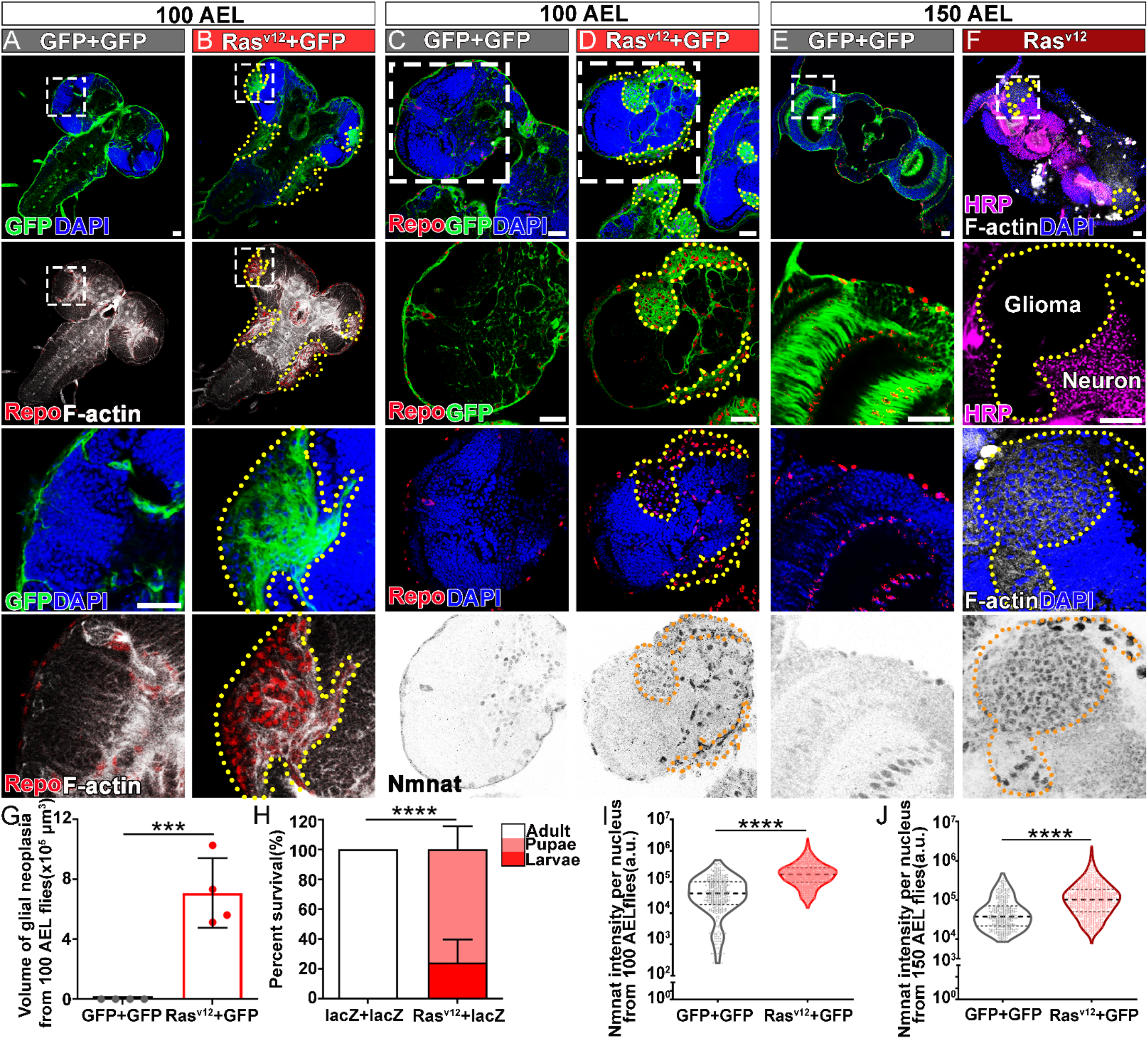
NMNAT is upregulated in *Ras^v12^*-induced glial neoplasia in *Drosophila*. (**A, B**) Larval CNS at 100 AEL with glial expression of GFP+GFP or Ras^v12^ +GFP was probed for F-actin (white), Repo (red), and DAPI (blue). The yellow dashed lines mark the boundary of glial neoplasia. The third and fourth rows show the boxed area of the first and second rows. (**C-F**) Larval CNS at 100 (**C, D**) and 150 (**E, F**) AEL. The second to forth rows show the boxed areas in the first row. (**C-E**) Brains were probed for Nmnat (grey), Repo (red), and DAPI (blue). (**F**) Brains were probed for HRP (magenta), Nmnat (grey), F-actin (white), and DAPI (blue). Yellow dashed lines mark the glial neoplasia boundaries. (**G**) Quantification of the total glial neoplasia volumes in each fly. Data are presented as mean ± s.d., n = 4. Significance level was established by one-way ANOVA post hoc Bonferroni test. (**H**) Survival rate. Data are presented as mean ± s.d., n ≥ 3. Significance level was established by Chi-square test. (**I-J**) Nmnat intensity at 100 AEL and 150 AEL. Data are presented as median ± quartiles, n ≥ 3. Significance level was established by one-way ANOVA post hoc Bonferroni test. \*\*\**P* ≤ 0.001. \*\*\*\**P* ≤ 0.0001. Scale bars, 30 µm.

### NMNAT is required for glial neoplasia development in *Drosophila*

To determine whether increased NMNAT is required for glial neoplasia development, we used the RNAi approach to downregulate NMNAT expression in *Ras^v12^*-induced glial neoplasia cells (Brazill, Cruz, Zhu, & Zhai, 2018). Interestingly, knocking down Nmnat drastically reduced both the volume and the number of individual glial cells in the brain and VNC at 100 hours AEL (Fig. 2A, C, D), demonstrating a strong antitumor effect of NMNAT inhibition *in vivo*. We analyzed RNAi-mediated knockdown of NMNAT in normal glial cells (without Ras^v12^ expression) did not result in growth inhibition (Figure 2-figure supplement 1), suggesting NMNAT is not essential for healthy cell survival. In addition, we found NMNAT expression level in NMNAT RNAi and Ras^v12^ overexpression flies is lower than wild-type flies (Figure 2-figure supplement 3).

**Figure 2 with 3 supplements.**
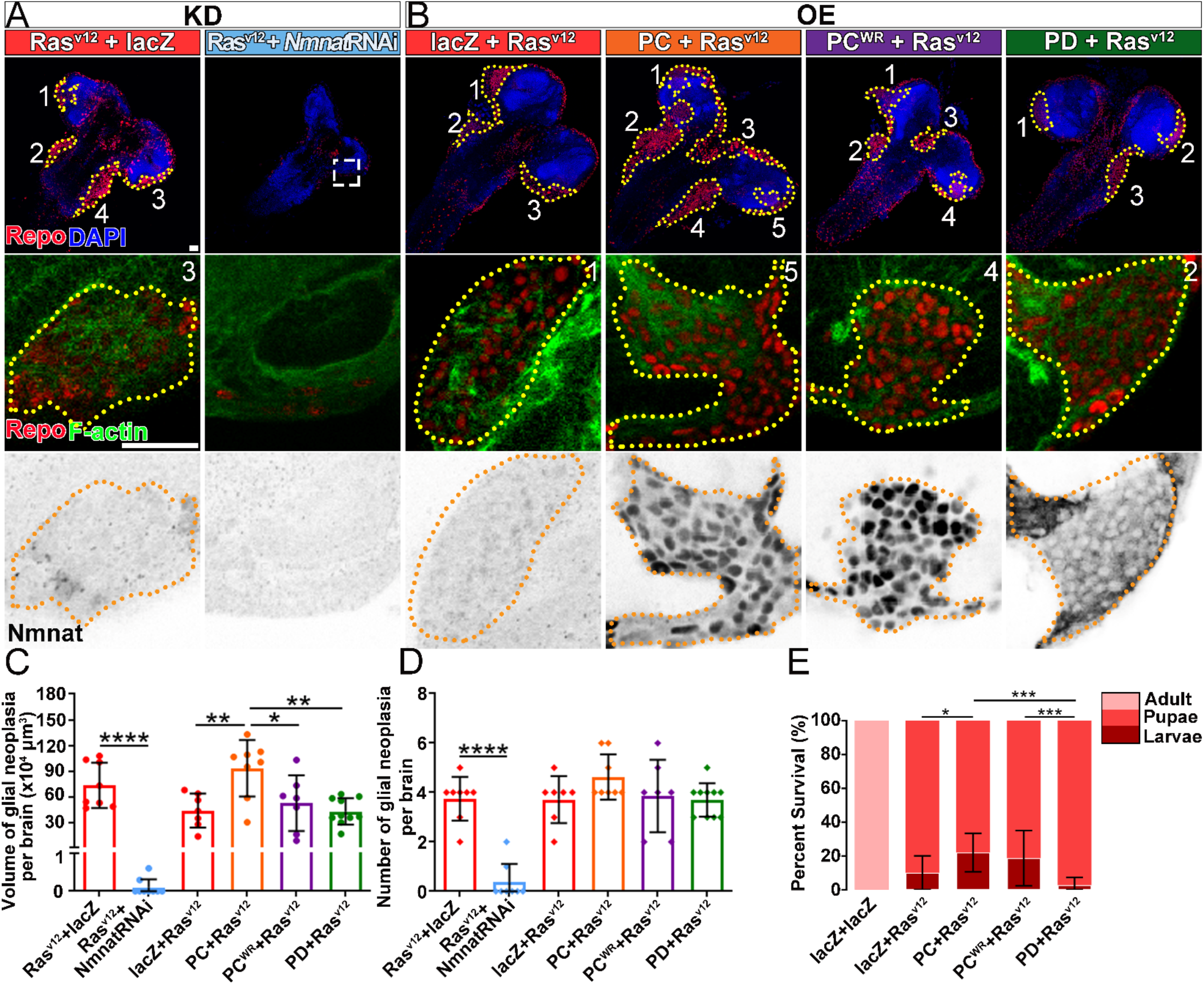
NMNAT is required for glial neoplasia growth in *D*rosophila. (**A, B**) Larval CNS at 100 AEL with glial expression of Ras^v12^ + lacZ, Ras^v12^ + NmnatRNAi, lacZ+Ras^v12^, PC+Ras^v12^, PC^WR^+Ras^v12^ and PD+Ras^v12^ was probed for F-actin (green), Repo (red), DAPI (blue), and Nmnat (grey). Each individual glial neoplasia is marked with dashed lines and numbered. The second and third rows show the high magnification of glial neoplasia areas in the first row. Scale bars, 30 µm. (**C**) Quantification of glial neoplasia volume in each fly. Data are presented as mean ± s.d., n ≥ 7. Significance level was established by one-way ANOVA post hoc Bonferroni test. (**D**) Quantification of glial neoplasia number in each fly. Data are presented as mean ± s.d., n ≥ 7. Significance level was established by one-way ANOVA post hoc Bonferroni test. (**E**) Survival rate of flies with glial expression of Ras^v12^ together with lacZ, PC, PC^WR^ or PD. Data are presented as mean ± s.d., n ≥ 3. Significance level was established by chi-square test. \**P* ≤ 0.05. \*\**P* ≤ 0.01. \*\*\**P* ≤ 0.001. \*\*\*\**P* ≤ 0.0001.

Next, we tested whether upregulating NMNAT can promote glial neoplasia formation and progression. *Drosophila* has one *Nmnat* gene, expressing two protein isoforms through alternative splicing, a nuclear isoform Nmnat-PC and a cytosolic isoform Nmnat-PD. The isoforms share similar enzymatic activity but have distinct chaperone properties. In addition, Nmnat-PC (nuclear) and Nmnat-PD (cytoplasmic) are differentially regulated under stress conditions (Ruan, Zhu, Li, Brazill, & Zhai, 2015). In *Ras^v12^*-induced glial neoplasia, dramatically increased Nmnat is mainly observed in the nuclear region (Fig. 1D and F), likely to be the Nmnat-PC (nuclear) isoform. To further evaluate the compartmentalized role of NMNAT during glial neoplasia formation, we generated flies expressing *Ras^v12^* together with Nmnat-PC (nuclear) or Nmnat-PD (cytoplasmic). As shown previously, Nmnat-PC (nuclear) is highly enriched in the nucleus and colocalizes with the nuclear marker Repo, while Nmnat-PD is predominantly cytoplasmic (Ruan et al., 2015). Interestingly, overexpression of Nmnat-PC (nuclear), but not Nmnat-PD (cytoplasmic), significantly increased the total volumes of glial neoplasia (Fig. 2B and C). We also analyzed the number of glial neoplasia and observed no significant difference among the groups (Fig. 2D). The lethality of the flies (Fig. 2E) was positively correlated with glial neoplasia size and overexpression of Nmnat-PC (nuclear) significantly increased the lethality.

To determine whether the enzyme activity of NMNAT is required for glial neoplasia tumorigenesis, we generated flies expressing an enzyme inactive mutant Nmnat-PC (nuclear) isoform (PC^WR^) where two key residues for substrate binding were mutated (Figure 2-figure supplement 2) (Zhai et al., 2006). We found that Nmnat-PC^WR^ (nuclear) overexpression did not significantly affect glial neoplasia volumes or numbers or survival outcome when compared to the control (Fig. 2C-E). These results suggest that nuclear enzymatically active NMNAT promoted glial neoplasia growth.

### NMNAT is essential to the proliferation of human glioma cells

We next examined the function of NMNAT in human glioma cell proliferation, specifically human NMNAT1 (nuclear) and NMNAT2 (cytoplasmic) (Berger et al., 2005). We determined NMNAT1 and NMNAT2 protein levels in human glioma cells and normal astroglia cells (SVG p12). Compared to SVG p12 cells, NMNAT1 and NMNAT2 are increased in both glioma cells T98G and U87MG (Figure 3-figure supplement 3). Then, we manipulated the expression of NMNAT by siRNA-mediated knockdown and plasmid-mediated overexpression in T98G cells and monitored real-time cell growth using the xCELLigence platform (Ke, Wang, Xu, & Abassi, 2011). Interestingly, we found T98G cell proliferation was drastically inhibited when NMNAT1 or NMNAT2 was downregulated (Fig. 3A). This observation was confirmed and extended in the MTT assay (van Meerloo, Kaspers, & Cloos, 2011), where cell proliferation was reduced in NMNAT1- or NMNAT2- knockdown cells (Figure 3-figure supplement 1). In contrast, overexpressing NMNAT1 or NMNAT2 promoted cell growth (Fig. 3D). Moreover, we used a plate colony formation assay to determine the tumorigenic potential of cells (Franken, Rodermond, Stap, Haveman, & van Bree, 2006). We found that knockdown of NMNAT1 or NMNAT2 reduced the colony number of T98G, while overexpression of NMNAT1 or NMNAT2 increased colony formation (Fig. 3B-F). These results are consistent with the genetic requirement of NMNAT observed in the fly glial neoplasia, suggesting the conservation of NMNAT function in promoting glioma cell growth and proliferation.

**Figure 3 with 3 supplements.**
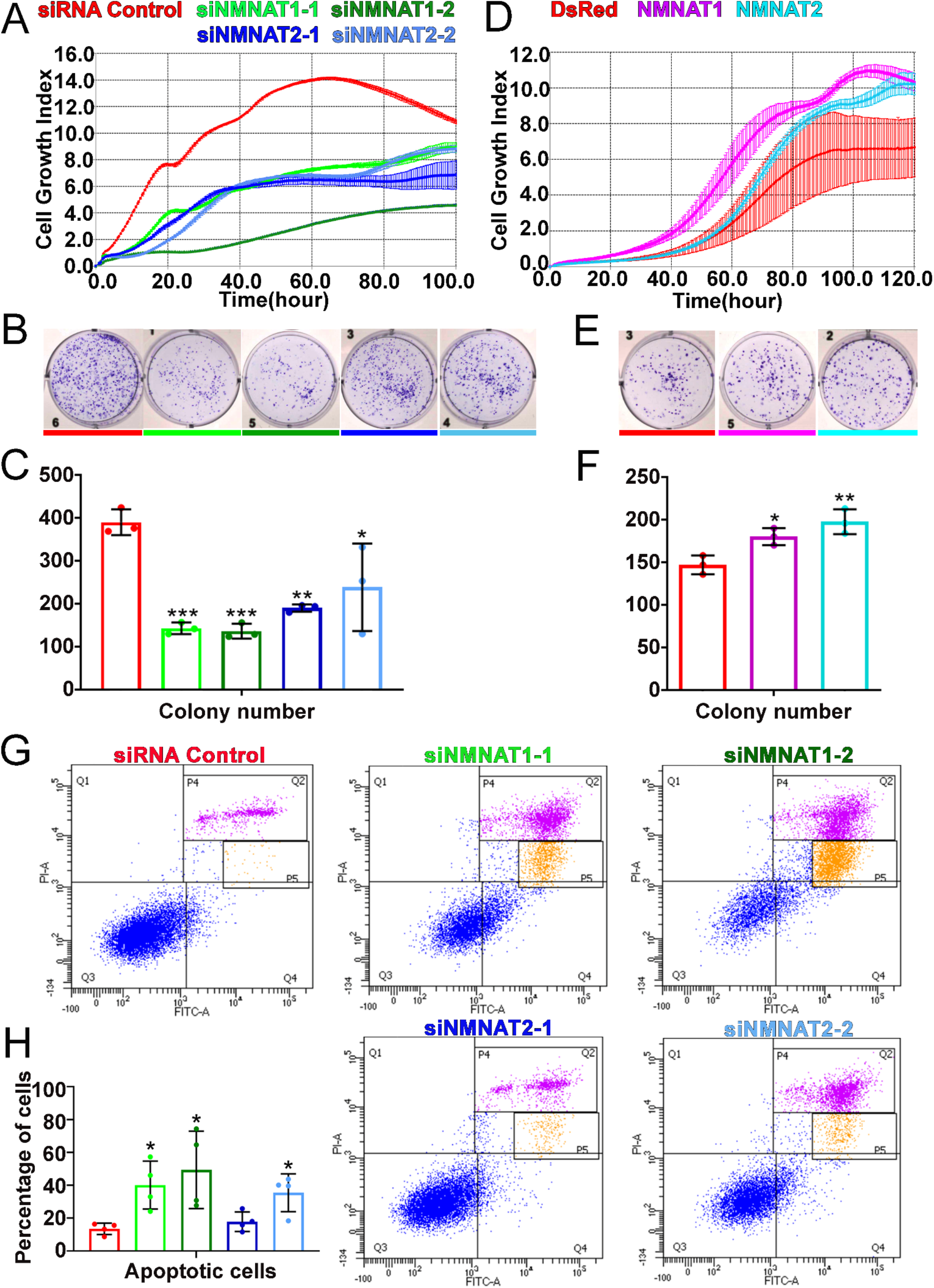
NMNAT expression is essential to the proliferation of human GBM cells. (**A, D**) The xCELLigence real-time cell analysis assay was used to monitor the growth index of T98G cells after NMNAT knockdown by transfecting siNMNAT1 or siNMNAT2, or after NMNAT overexpression by transfecting NMNAT1 or NMNAT2 plasmid. Cells transfected with siRNA control or DsRed were used as controls. (**B, E**) Colony formation assay was used to measure the colony formation capabilities of T98G cells after NMNAT knockdown by transfecting siNMNAT1 or siNMNAT2, or after NMNAT overexpression by transfecting NMNAT1 or NMNAT2. Cells transfected with siRNA control or DsRed were used as controls. (**C, F**) Quantification of the colony number in (**B, E**). Data are presented as mean ± s.d. n = 3. Significance level was established by one-way ANOVA post hoc Bonferroni test. (**G**) T98G cell apoptosis was detected by flow cytometry after NMNAT knockdown. P4 and P5 apoptotic populations were shown with magenta and yellow separately. (**H**) Quantification of apoptotic cells rate of siRNA control, siNMNAT1-1, siNMNAT1-2, siNMNAT2-1 and siNMNAT2-2. Q2 and Q4 was quantified as apoptotic cells. Data are presented as mean ± s.d. n = 4. Significance level was established by t-test. \**P* ≤ 0.05. \*\**P* ≤ 0.01. \*\*\**P* ≤ 0.001.

To further determine whether NMNAT is involved in apoptosis of glioma, we carried out an apoptosis detection assay through flow cytometry in T98G cells. We transfected siRNA targeting NMNAT into T98G cells and then analyzed Annexin V-FITC/PI by flow cytometric 72 hr post transfection. Interestingly, we found that knockdown of NMNAT, at the knockdown rate of 40-50% for NMNAT1 or at 20-30% for NMNAT2, significantly increased the percentage of apoptotic cells, including early apoptotic and late apoptotic cells (Fig. 3G and H). We also examined the cell cycle distribution of these cells. The cell cycle assay showed G2/M phase was only slightly increased in T98G cells with NMNAT1 knockdown (Figure 3-figure supplement 2). These results suggest NMNAT promotes glioma cell growing mainly through inhibiting cell apoptosis.

### Overexpression of NMNAT decreases caspase-3 activation in glioma

The cysteine-dependent proteases (caspases) are activated by upstream proteins to mediate apoptosis (Kurokawa & Kornbluth, 2009). Caspase-3 is the main effector protease cleaving a large number of substrates during apoptosis. Previous studies revealed that nuclear translocation and accumulation of caspase-3 play a critical role in the progression of apoptosis (Prokhorova, Kopeina, Lavrik, & Zhivotovsky, 2018). Caspase-mediated pathway is highly conserved in mammalian and *Drosophila* (Fuchs & Steller, 2011; Shi, 2001) (Figure 4-figure supplement 1A). To validate the role of caspase pathway in *Drosophila* glial neoplasia, we examined tumor growth in flies with downregulation of DCP1, the homolog of caspase-3/7 in mammalian. In these flies, glial neoplasia volume was significantly increased (Figure 4-figure supplement 1B and C), suggesting the important role of caspase-mediated apoptosis in *Drosophila* glial neoplasia progression. To test whether NMNAT regulates this process, we determined the localization and protein levels of caspase-3 in the glial neoplasia with overexpression of different Nmnat isoforms. We used Repo and DAPI to label the nuclei region and observed a significant decrease of caspase-3 level in glial neoplasia that overexpressing Nmnat-PC (nuclear), compared with those overexpressing lacZ, Nmnat-PC^WR^ (nuclear), or Nmnat-PD (cytoplasmic) (Fig. 4A and C). In addition, when we knocked down Nmnat in Ras^v12^ expressing glial cells, we observed significant nuclear enrichment of caspase-3 (Fig. 4B and D). These results suggest that NMNAT is a negative regulator of glial neoplasia cell apoptosis in *Drosophila*.

**Figure 4 with 1 supplement.**
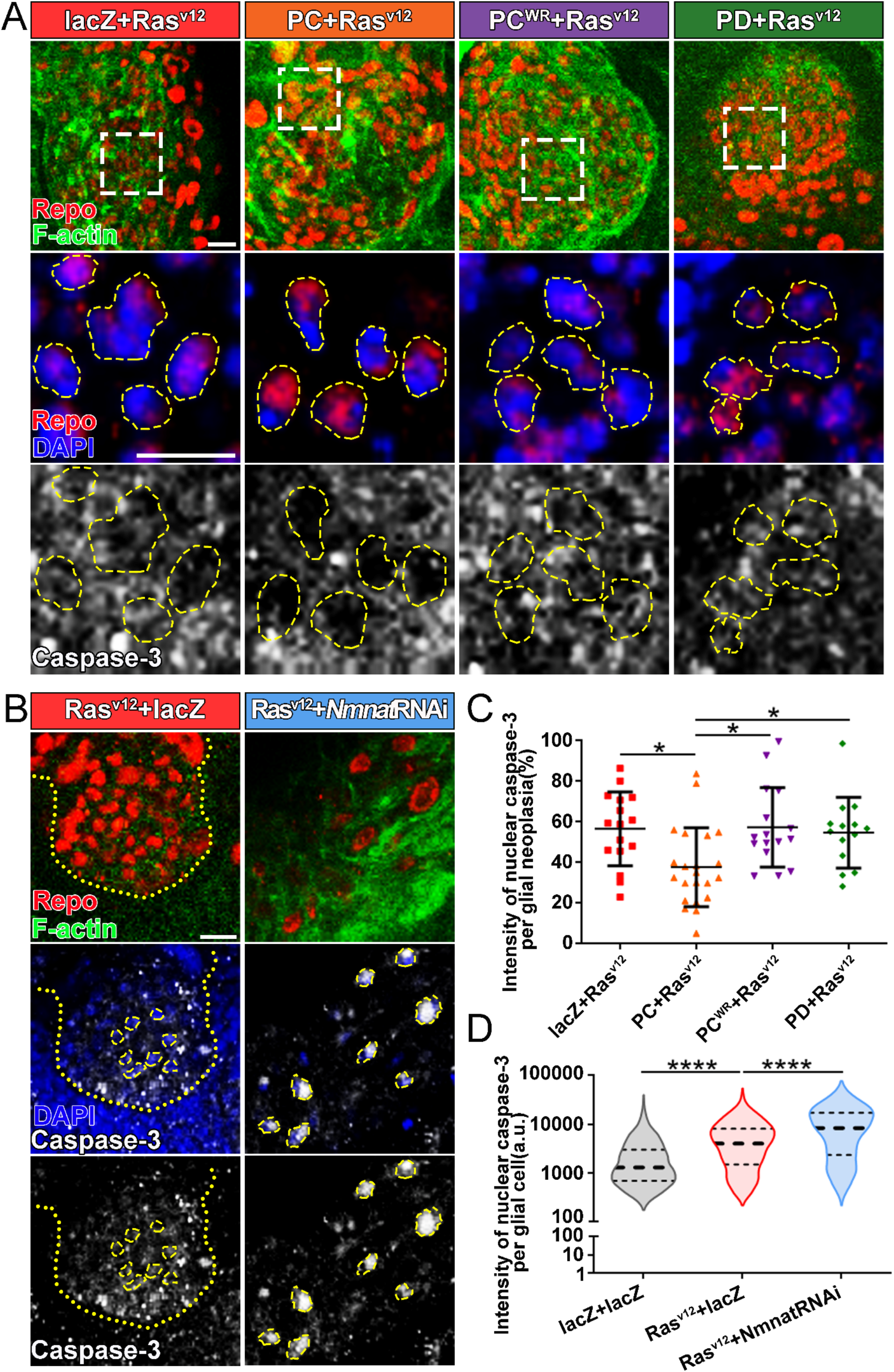
Overexpression of NMNAT decreases caspase-3 activation in glial neoplasia. (**A**) Glial neoplasia from files expressing lacZ, PC, PC^WR^ or PD were probed for Repo (red), F-actin (green), DAPI (blue), and caspase-3 (grey). The top row shows the whole glial neoplasia area. The second and third rows are the high magnification of the boxed areas in the first row. Yellow dashed lines indicate the nuclear area. (**B**) Glial neoplasia from flies expressing lacZ or Nmnat RNAi were probed for Repo (red), F-actin (green), DAPI (blue), and caspase-3 (grey). Yellow dot lines indicate glial neoplasia boundary in the Ras^v12^ + lacZ group. Yellow dashed lines indicate the boundaries of the nucleus and cytoplasm. Scale bars, 10 µm. (**C**) Quantification of the percentage of nuclear caspase-3 intensity per glial neoplasia. Data are presented as mean ± s.d. n ≥ 3. Significance level was established by one-way ANOVA post hoc Bonferroni test. (**D**) Quantification of the nuclear caspase-3 per glial cell. Data are presented as median ± quartiles, n ≥ 3. Significance level was established by one-way ANOVA post hoc Bonferroni test. \**P* ≤ 0.05. \*\*\*\**P* ≤ 0.0001.

Next, we examined apoptosis and the activation of caspase-3 in human T98G cells. We found that knockdown of NMNAT led to increased nuclear caspase-3 (Fig. 5A and C). Western blot analysis showed a specific increase of fully processed P17/19 species of cleaved caspase-3 (Fig. 5D), indicating the activation of apoptosis (Porter & Janicke, 1999). To examine the effect of overexpressing NMNAT on apoptosis, we employed cisplatin treatment to induce apoptosis as the basal level of apoptosis in T98G glioma cells is low (Kondo et al., 1995). Cisplatin significantly increased nuclear caspase-3 levels as expected. Interestingly, overexpression of either NMNAT1 or NMNAT2 reduced nuclear caspase-3 in cisplatin-induced apoptosis (Fig. 5B and E), specifically the fully processed cleaved caspase species P17/19 as shown by western blot analysis (Fig. 5F). Taken together, these results suggest that NMNAT promotes glioma growth by inhibiting caspase-mediated apoptosis.

**Figure 5 with 2 supplements.**
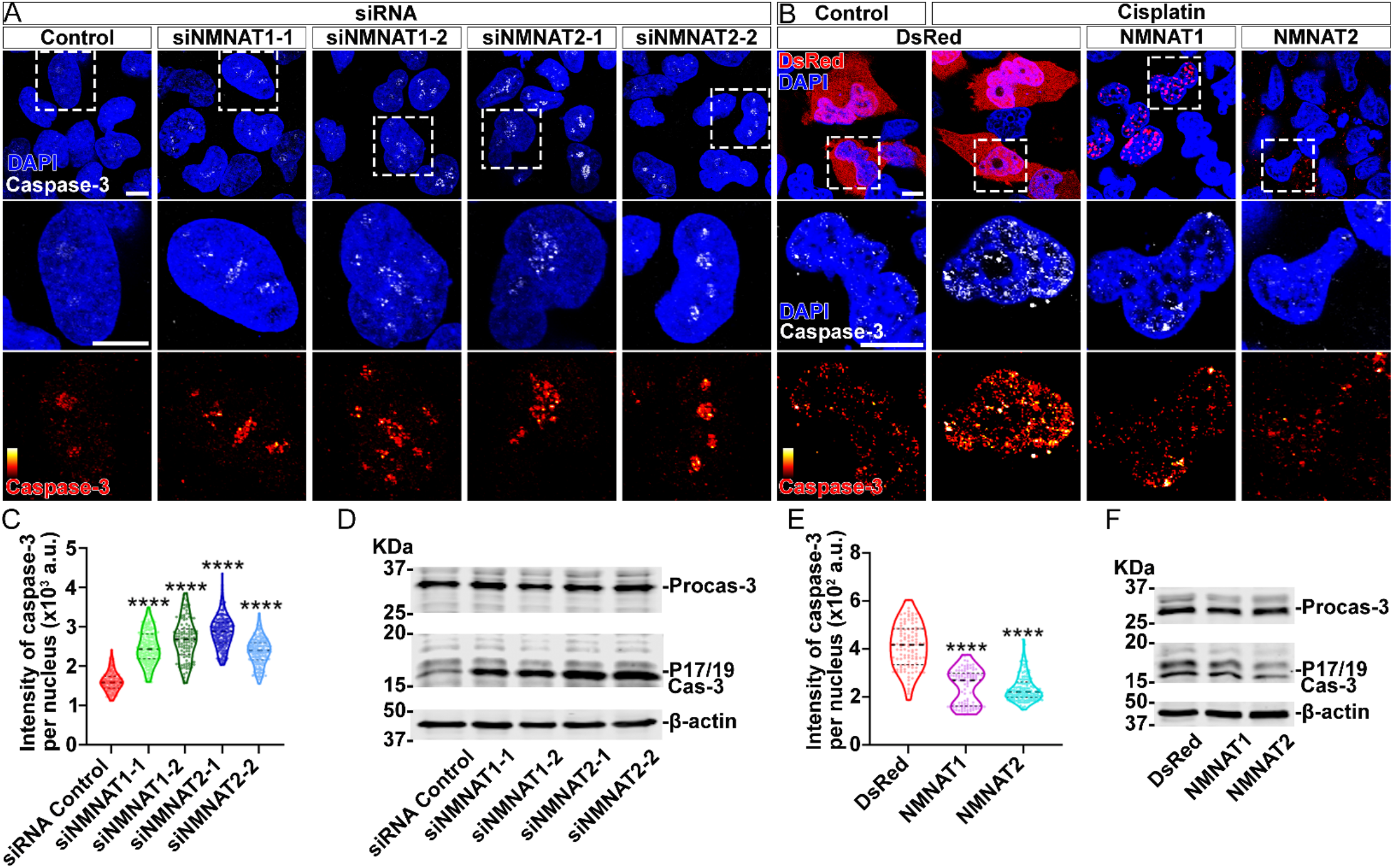
NMNAT decreases caspase-3 activation in human glioma cells. (**A**) T98G cells were transfected with siNMNAT1 or siNMNAT2 and stained with DAPI (blue) and caspase-3 (white). (**B**) T98G cells were transfected with DsRed (red), DsRed-NMNAT1 (red), or DsRed-NMNAT2 (red), treated with cisplatin 8 hours after transfection, and stained with DAPI (blue) and caspase-3 (grey). The second and third rows are the high magnification of the boxed areas in the first row. In the third row, the intensity of caspase-3 is indicated by a heat map (0-4095). Scale bars, 10 µm. (**C**) Quantification of nuclear caspase-3 intensity in A. Data are presented as Median ± quartiles, n ≥ 100. Significance level was established by one-way ANOVA post hoc Bonferroni test. (**E**) Quantification of nuclear caspase-3 intensity in B. Data are presented as Median ± quartiles, n ≥ 100. Significance level was established by one-way ANOVA post hoc Bonferroni test. (**D, F**) Proteins were extracted from T98G cells transfected with siRNA (**D**), plasmids and treated with cisplatin 8 hours (**F**) for western blot analysis. P17/19 was considered as cleaved caspase-3. β-actin was used as an internal control. \*\*\*\**P* ≤ 0.0001.

### Overexpression of NMNAT increases DNA damage tolerance and decreases nuclear p53 in glial neoplasia

Increased DNA damage is one of the hallmarks of cancer. Two common strategies cancer cells use to avoid the triggering of cell apoptosis by DNA damage are hyperactivating DNA damage repair, and inactivating cell apoptosis initiation (Norbury & Zhivotovsky, 2004). Since NAD^+^ plays important regulatory roles in both DNA damage repair and cell apoptosis, and NAD^+^ synthase activity is required for glial neoplasia growth (Fig. 2), we next examined the effect of NMNAT on the DNA damage pathway in glioma. We first determined DNA damage by using a phosphor-specific antibody to histone 2A variant (H2Av), a marker for DNA double-strand breaks (Lake, Holsclaw, Bellendir, Sekelsky, & Hawley, 2013). We observed a significant elevation of H2Av signal in Nmnat- PC (nuclear) overexpressing brains compared to that in Nmnat-PD (cytoplasmic), Nmnat-PC^WR^ (nuclear), or lacZ overexpressing brains (Fig. 6A), suggesting DNA damage level is higher in glial neoplasia with Nmnat-PC (nuclear) overexpression.

**Figure 6.**
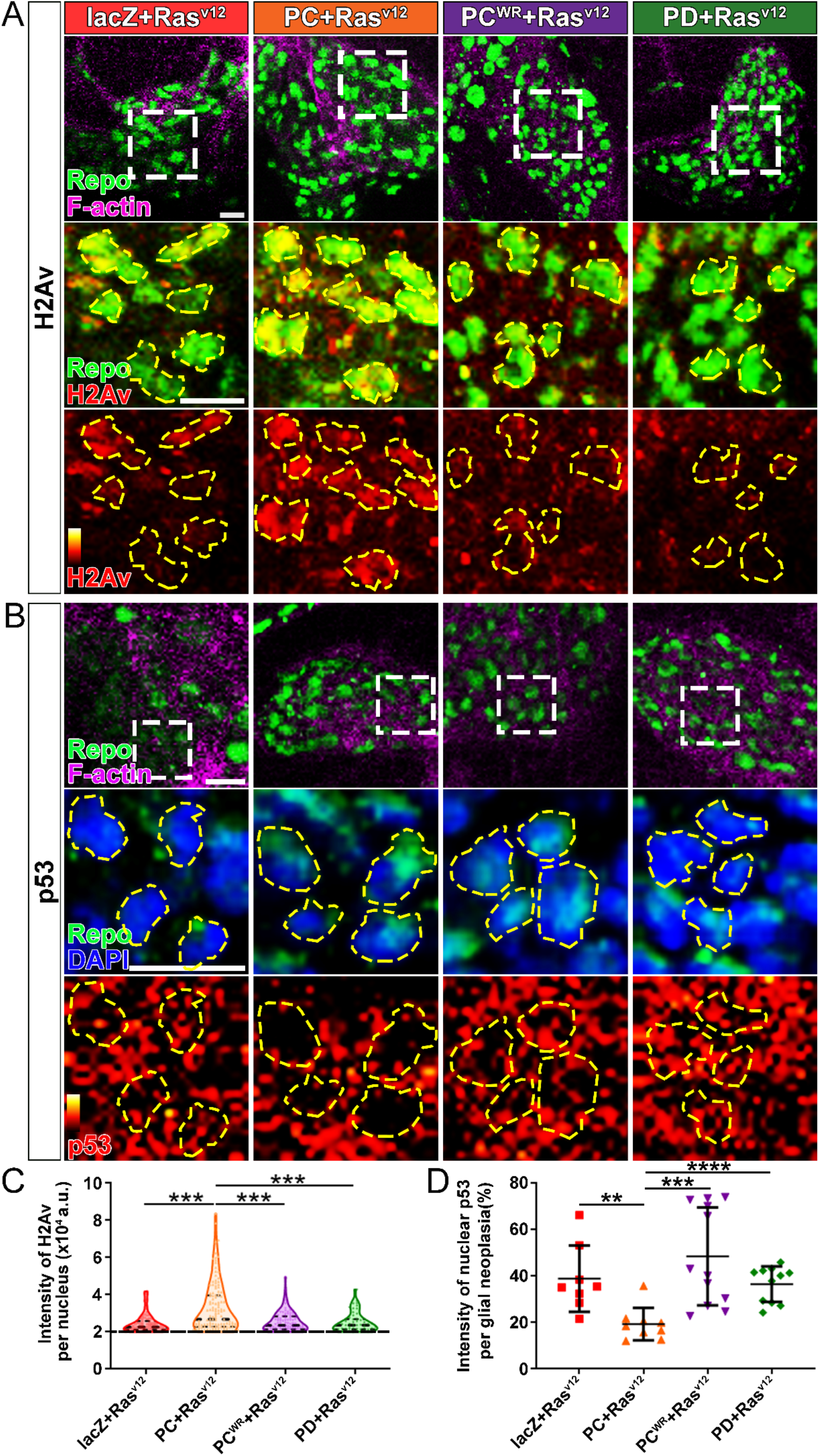
Nmnat-PC inhibits DNA damage-induced p53 activation in glial neoplasia. (**A**) Glial neoplasia from flies expressing lacZ, PC, PC^WR^ or PD were stained with H2Av (red), Repo (green), and F-actin (magenta). The second and third rows are high magnification of the boxed areas in the first row. In the third row, the intensity of H2Av is indicated by a heatmap (0-4095). (**B**) Glial neoplasia from flies expressing lacZ, PC, PC^WR^ or PD were stained with p53, Repo (green), F-actin (magenta), and DAPI (blue). The second and third rows are high magnification of the boxed areas in the first row. In the third row, the intensity of p53 is indicated by a heatmap (0-4095). Yellow dashed lines indicate the nuclear areas. Scale bars, 10 µm. (**C**) Quantification of H2Av intensity in Repo-positive cells. The black dashed line indicates the threshold. According to the lacZ group, value 20,000 is set as the threshold. Data are presented as median ± quartiles, n ≥ 3. Significance level was established by one-way ANOVA post hoc Bonferroni test. (**D**) Quantification of nuclear p53 intensity. Data are presented as mean ± s.d., n ≥ 3. Significance level was established by t-test. \*\**P* ≤ 0.01. \*\*\**P* ≤ 0.001. \*\*\*\**P* ≤ 0.0001.

We next examined the distribution of endogenous p53 in glial neoplasia and found that while in control glial neoplasia cells (LacZ group), p53 was relatively evenly distributed with ∼40% of p53 in the nucleus, a significantly reduced nuclear p53 pool (∼20%) was found in Nmnat-PC (nuclear) overexpressing glial neoplasia cells (Fig. 6B and D). Together with the observation of higher levels of DNA damage in Nmnat-PC (nuclear) overexpressing glial neoplasia cells, these results indicate that Nmnat-PC (nuclear) expression potentially regulates p53 response to DNA damage to allow higher tolerance to DNA damage.

p53 is well known as a key player controlling cell fate in response to DNA damage: initiate DNA repair when there is limited DNA damage, and induce apoptosis when DNA damage is too severe (Roos & Kaina, 2013). To validate the role of p53 in glial neoplasia development in *Drosophila*, we examined the effect of a p53 inhibitor: Pifithrin-α (PFT-α). PFT-α is reported to inhibit translocation of p53 and affect p53 related transactivation (Komarov et al., 1999; Leker, Aharonowiz, Greig, & Ovadia, 2004; Murphy et al., 2004). We analyzed glial neoplasia tissue volume with GFP and DAPI staining in the central nervous system (CNS) of flies (Fig. 7A). The glial neoplasia volume was significantly increased in PFT-α -treated flies compared to that in DMSO-treated flies (Fig. 7C). The increase in glial neoplasia volume was accompanied by a decrease in survival (Fig. 7B), and the reduced cleaved caspase-3 intensity (Fig. 7D). These results suggest p53 is critical for glial neoplasia progression in *Drosophila*, and p53 inhibition phenocopies NMNAT overexpression in glial neoplasia growth.

**Figure 7.**
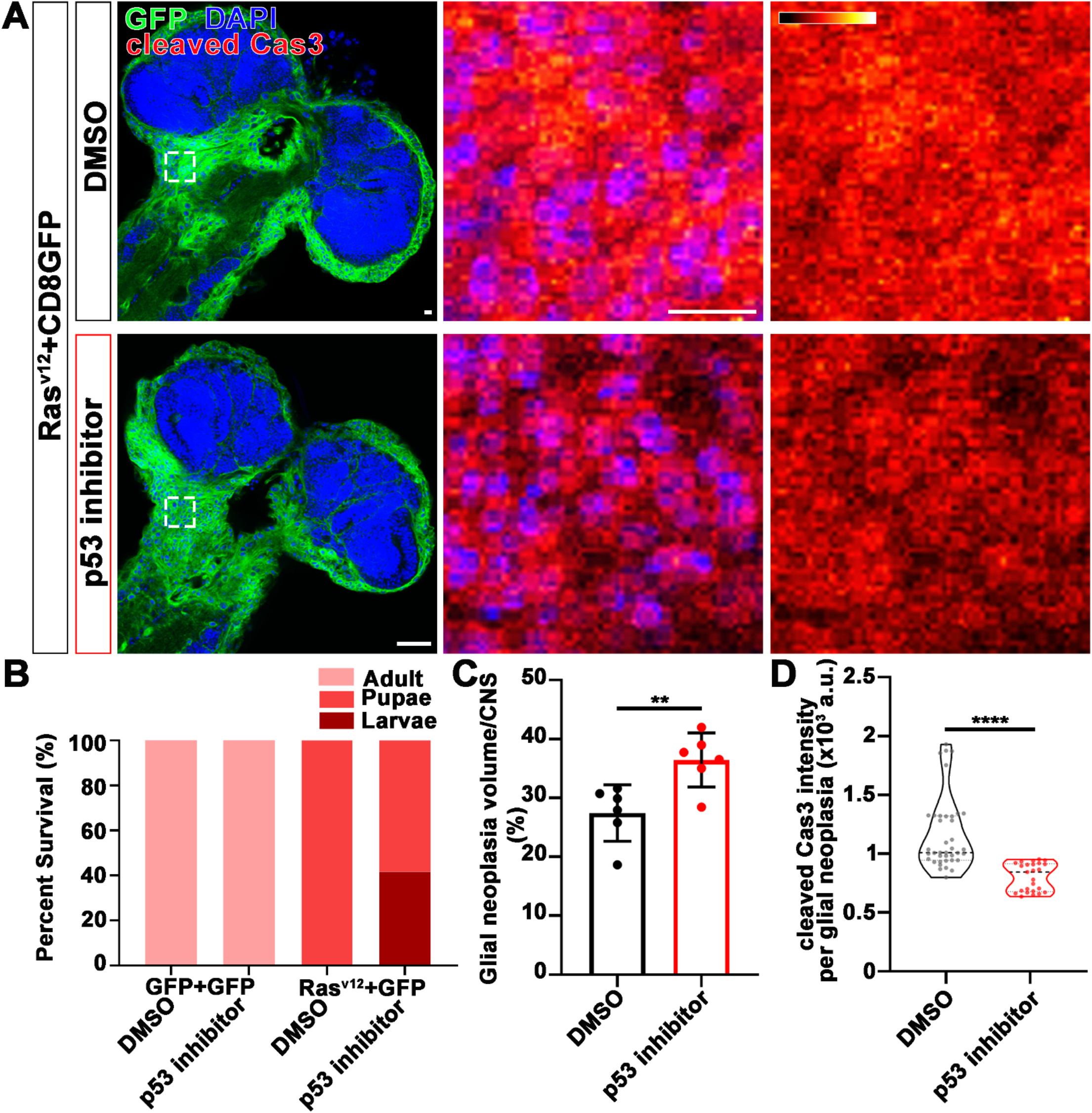
p53 inhibitor increases glial neoplasia volume in CNS and larvae lethality. (**A**) Flies expressing Ras^v12^ and CD8GFP were treated with DMSO or p53 inhibitor respectively, stained with Cleaved Caspase-3 (red) and DAPI (blue). The first column is the whole CNS of flies. White dashed lines indicate the glial neoplasia areas. The second and third columns are high magnification of the boxed white areas in the first row. The intensity of cleaved caspase-3 is indicated by a heatmap (0-4095). Scale bars, 10 µm. (**B**) Survival rate of flies. (**C**) Quantification of ratio of glial neoplasia volumes in CNS. Data are presented as mean ± s.d., n ≥ 3. Significance level was established by one-way ANOVA post hoc Bonferroni test. (**D**) Quantification of cleaved caspase-3 intensity. Data are presented as median ± quartiles, n ≥ 3. Significance level was established by one-way ANOVA post hoc Bonferroni test. \*\**P* ≤ 0.01. \*\*\*\**P* ≤ 0.0001.

### NMNAT regulates PARylation and acetylation of p53

Our observations that NMNAT overexpression-induced higher tolerance to DNA damage and altered p53 response is intriguing. Maintaining functional DNA repair is critical for cancer cells to survive during rapid cell proliferation and constant replication of DNA. In response to DNA damage, NAD^+^-dependent PARP1 PARylates a large number of proteins, including p53, which is the largest NAD^+^ consumer in the nucleus (Kim, Zhang, & Kraus, 2005). It has been shown that PARylated p53 has reduced stability and activity (Simbulan-Rosenthal, Rosenthal, Luo, & Smulson, 1999). We hypothesize that NMNAT regulates PARylation in glioma. To test this hypothesis, we first examined the level of protein PARylation under the conditions NMNAT overexpression, and found that protein PARylation level was significantly increased with NMNAT1 or NMNAT2 overexpression and significantly reduced with siRNA knockdown using dot blot analysis (Fig. 8A, B and Figure 8-figure supplement 1). Next, we examined the protein-protein interaction among p53, NMNAT1 and PARP1 using immunoprecipitation. Interestingly, we detected PARP1 and NMNAT proteins in the p53-immunoprecipitated fraction (Fig. 8C). Furthermore, although the level of total p53 was not significantly affected by NMNAT expression, the level of PARP1 immunoprecipitated with p53 was increased with the overexpression of NMNAT (Fig. 8C). These results suggest the presence of a trimeric complex of p53, NMNAT, and PARP1, and the potential role of NMNAT in promoting the trimeric complex formation.

**Figure 8 with 1 supplement.**
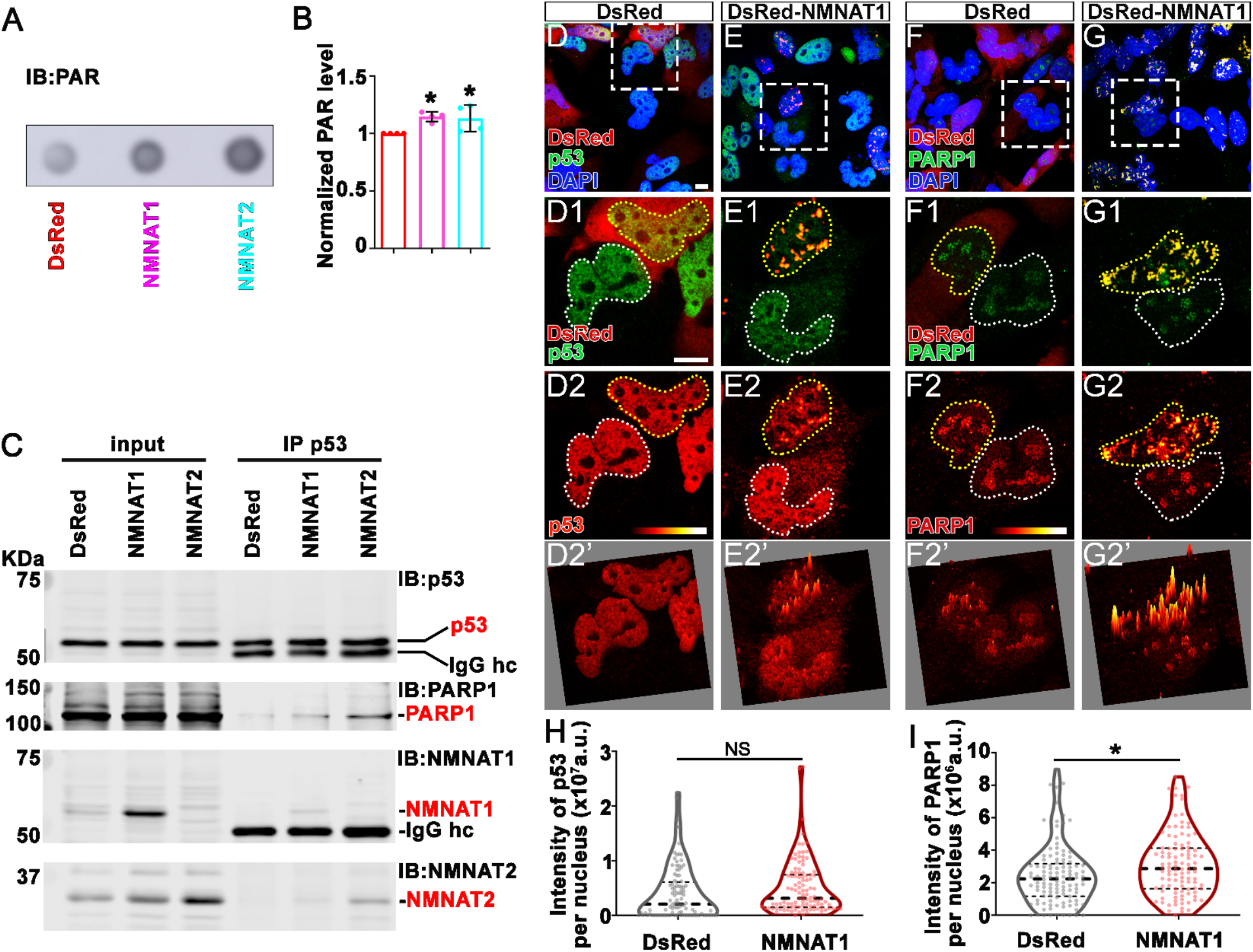
NMNAT interacts with PARP1 and upregulates PARylation of p53. (**A**) Proteins were extracted from T98G cells transfected with plasmids for dot blot analysis using anti-PAR antibody. (**B**) Quantification of dot blot normalized with β-actin as internal control. Data are presented as mean ± s.d., n = 4. Significance level was established by t-test. (**C**) Protein samples extracted from T98G cells transfected with DsRed, DsRed-NMNAT1 or NMNAT2 were immunoprecipitated (IP) with a p53 antibody and subjected to immunoblot (IB) analysis for p53, PAR, PARP1, NMNAT1 and NMNAT2. (**D-G**) T98G cells transfected with DsRed or DsRed-NMNAT1 were stained for DAPI (blue), p53 (green), or PARP1 (green). The second to the fourth rows are high magnification of the boxed area in the first row. The intensity (0-4095) of p53 or PARP is indicated in a heat map (**D2-G2**) or surface plot (**D2’-G2’**). Scale bars, 10 µm. (**H**) Quantification of nuclear p53. Data are presented as median ± quartiles, n ≥ 100. Significance level was established by one-way ANOVA post hoc Bonferroni test. (**I**) Quantification of PARP1 intensity. Data are presented as median ± quartiles, n ≥ 100. Significance level was established by one-way ANOVA post hoc Bonferroni test. \**P* ≤ 0.05. NS, not significant.

To confirm and extend the biochemical analysis, we carried out immunofluorescent colocalization studies of T98G glioma cells expressing NMNAT1 and detected the colocalization of NMNAT1 and p53 (Fig. 8E1) and NMNAT1 and PARP1 (Fig. 8G1). Consistent with western analysis (Fig. 8C), p53 protein level is not altered by NMNAT expression as p53 immunofluorescence intensity was similar between NMNAT1 expression cells and neighboring desired expressing control cells (Fig. 8D2 and quantified in Fig. 8H). Interestingly, the distribution of p53 changed from a diffuse pattern to clustering to NMNAT1 positive hotspots, as visualized by fluorescence surface plot in Fig. 8E2’. Similarly, PARP1 protein also clustered to NMNAT1 positive hotspots (Fig. 8G2 and 8G2’), suggesting the close proximity of NMNAT, p53, and PARP1. In addition, in NMNAT expression cells, PARP1 level is slightly upregulated (Fig. 8I). Collectively, these results suggest NMNAT regulates p53 modification by complexing with p53 and PARP1, which may locally supply NAD^+^ to promote PARylation with high efficiency.

In addition to PARylation, another NAD^+^-dependent posttranslational modification of p53 is deacetylation. p53 is acetylated by p300/CBP and deacetylated by SIRTs family of NAD^+^-dependent deacetylases (Vaziri et al., 2001). SIRT1 is the major deacetylase regulating p53 activity through deacetylation of p53 at K382, and hence inhibiting the p53-mediated apoptosis pathway (Cheng et al., 2003). NMNAT1 has been reported to interact with SIRT1 directly (Zhang et al., 2009). We determined the level of acetyl-p53, by immunoprecipitating p53 from T98G glioma cells with or without NMNAT overexpression, and then probing for acetyl-p53 at K382. Interestingly, with NMNAT1 or NMNAT2 overexpression, acetyl-p53 was specifically reduced while total p53 levels remained the same (Fig. 9A and B), although a stable complex of p53 and SIRT1 was not detected. It is interesting to note that endogenous SIRT1 expression was upregulated in NMNAT overexpressing cells (Fig. 9C), suggesting a potential coregulation of NMNAT and SIRT1. To expand our analysis, we also repeated experiments in another human glioma cell line U87MG and observed consistent results (Figure 9-figure supplement 1). Collectively, these results show that NMNAT upregulation promotes the NAD^+^-dependent deacetylation of p53 and specifically reduces the pool of acetyl-p53.

**Figure 9 with 1 supplement.**
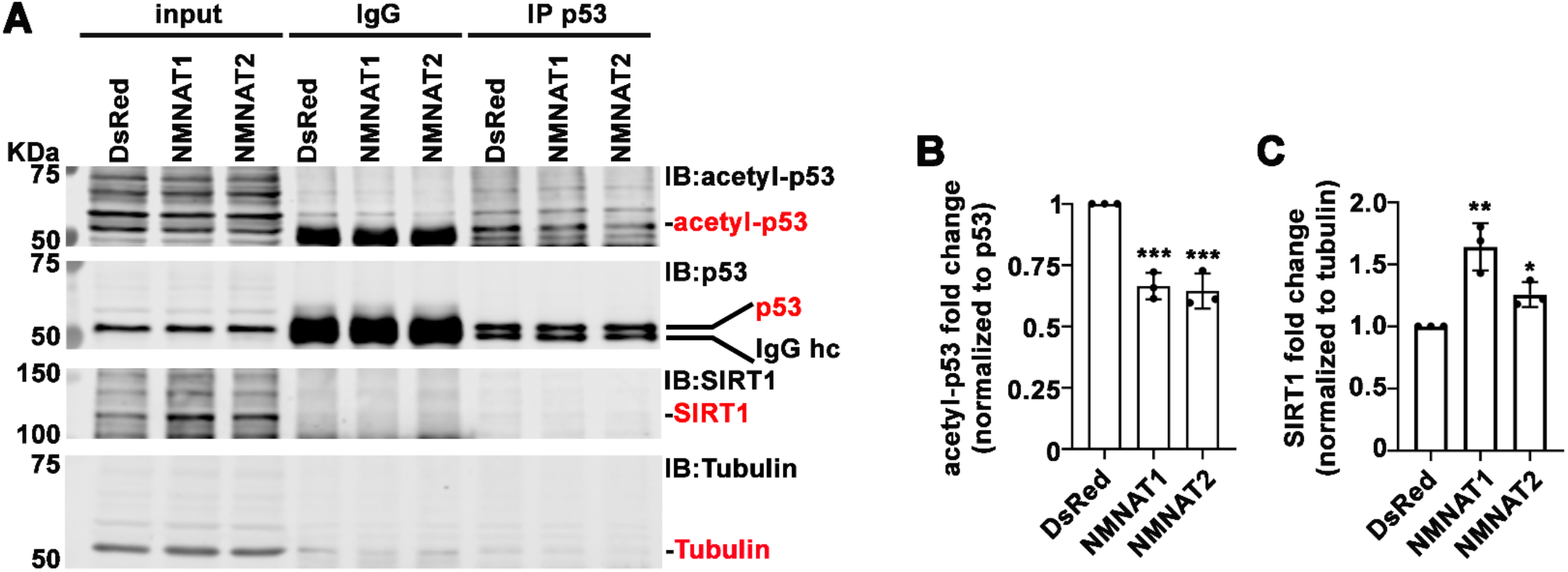
NMNAT upregulates SIRT1 and reduces acetylation of p53. **(A)** Protein samples extracted from T98G cells transfected with DsRed, DsRed-NMNAT1 or NMNAT2 were immunoprecipitated (IP) with a p53 antibody and probed for acetyl-p53 and SIRT1. (**B, C**) Quantification of acetyl- p53 and SIRT1. Data are presented as mean ± s.d., n = 3. Significance level was established by one-way ANOVA post hoc Bonferroni test. \**P* ≤ 0.05. \*\**P* ≤ 0.01. \*\*\**P* ≤ 0.001.

As both PARylation and deacetylation modifications of p53 have been reported to inactivate p53-mediate apoptosis induction (Juan et al., 2000; Luo, Su, Chen, Shiloh, & Gu, 2000; Malanga, Pleschke, Kleczkowska, & Althaus, 1998; Simbulan-Rosenthal et al., 1999), our results suggest

NMNAT promote glioma growth through facilitating NAD^+^-dependent post-translational modifications of p53 to ameliorate DNA damage-triggered apoptosis.

## DISCUSSION

In this study, we identified a critical role of NMNAT in promoting glioma cell proliferation and growth in a model of *Drosophila* glial neoplasia and human glioma cell lines. We found that NMNAT promotes glioma growth by allowing higher tolerance to DNA damage and inhibiting p53/caspase-mediated apoptosis. Mechanistically, upregulation of enzymatically active NMNAT promotes the NAD^+^-dependent post-translational modifications of p53, and specifically increases the PARylation of p53 and reduces the acetylation of p53. Furthermore, we detected a p53-NMNAT-PARP1 trimeric complex and increased SIRT1, suggesting a highly efficient NAD^+^-dependent post-translational modification process facilitated by NAD^+^ synthase NMNAT. Our findings support a tumorigenesis model where NMNAT proteins promote glioma growth through regulating NAD^+^-dependent post-translational modification of p53, and driving cellular pools of p53 toward PARylated-p53 (inactive p53) and away from acetyl-p53 (active p53) to ameliorate DNA damage-triggered cell death (Fig. 10).

**Figure 10 with 2 supplements.**
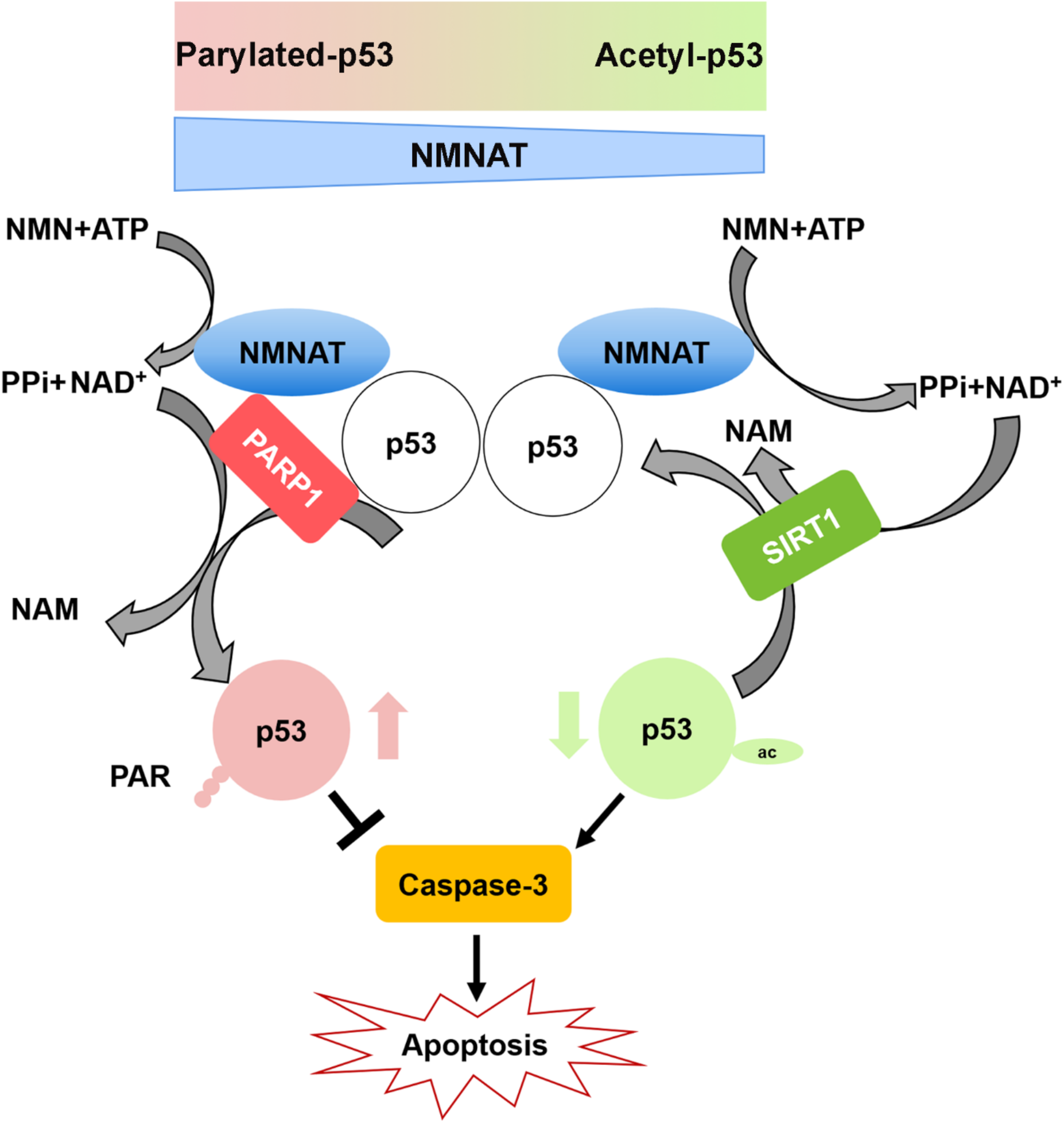
Diagram of cellular apoptosis response regulated by overexpressing NMNAT in glioma. In glioma cells, PAPR1 inhibits p53 activity by NAD^+^ dependent-poly(ADP-ribosyl)ation of p53 during DNA damage repair. NMNAT overexpression replenishes the NAD^+^ pool to promotes poly(ADP-ribosyl)ation and deacetylation of p53, suppressing p53 induced apoptosis, thereby leading to glioma growth.

### The advantages and potential of an *in vivo Drosophila* glial neoplasia

We adapted a glial neoplasia in *Drosophila* using the *UAS*-*Ras85D^v12^* and repo-GAL4 driver system that induces overgrowth of glial cells to mimic glial neoplasia formation (Read et al., 2009). Although RAS alterations in human glioma occur in lower frequency than some high alteration genes(Brennan et al., 2013), our rationale for using mutant RAS overexpressing model in *Drosophila* was to probe the shared (rather than Ras-specific) fundamental mechanisms in glial neoplasia. It would be an important future direction to establish *Drosophila* models using other high-frequency glioma drivers. Since all *Drosophila* glia express Repo, we can easily monitor the formation of *Ras^v12^*-driven glial neoplasia in the brain by GFP reporter, Repo, and F-actin labeling. In fluorescence imaging, normal brains typically have two to three layers of Repo-positive cells visible in each section (Fig. 1B). Therefore, any tissue mass consists of more than three layers of glia would be atypical and potentially tumor-like. We analyzed glial neoplasia with three key criteria: cell type (Repo-positive), cell number (more than three layers with at least 400), and tissue size (volume of at least 12.4 x 10^3^ μm^3^). Combined with our high-resolution imaging capability, these criteria allow us to distinguish tumor from non-glial neoplasia tissue with high confidence and to analyze glial neoplasia in the most robust and reproducible manner. In *Ras^v12^* expressing flies, we observed glial neoplasia occurred extensively in the brain and VNC.

In addition to the morphological phenotypes, we found that glial neoplasia reduced the animal survival rate. Specifically, the total volume of glial neoplasia tissue is positively correlated with the severity of reduced animal survival rate. Such correlation allows the use of high-resolution *in vivo* morphological imaging as a strong predictor of pathological outcome and a powerful tool to identify genetic modulators of tumorigenesis as we have done in this study, and potential pharmacological modulators for cancer therapy in the future.

### NMNAT-mediated NAD^+^ biosynthesis promotes glioma growth

Our results show that NMNAT expression promotes glioma progression but is likely dispensable for its initiation as NMNAT overexpression alone did not trigger tumorigenesis. Our results showed that the enzymatic function of NMNAT is required for glioma growth. This finding is not surprising given the fundamental role of NAD^+^ as a signaling cofactor that regulates cancer metabolism through its coenzymatic function in redox reactions underlying essential bioenergetic pathways including glycolysis, the tricarboxylic acid (TCA) cycle, and oxidative phosphorylation (Hanahan & Weinberg, 2000). While NAMPT is a uni-directional enzyme, synthesizing NMN (nicotinamide mononucleotide), NMNAT is downstream of NAMPT and directly regulates the level of NAD+ by catalyzing the reversible reaction of NAD^+^ synthesis. The direction of the reaction, forward (NAD+ production) or reverse (NAD+ breakdown), is dependent upon the availability of subtracts. Therefore, NMNAT functions as a cellular metabolic sensor and maintains the homeostasis of NAD^+^ pools.

The distribution of NAD+ is highly compartmentalized, with each subcellular NAD^+^ pool differentially regulated and preferentially involved in distinct NAD^+^-dependent signaling or metabolic events (Zhu, Liu, Park, Rai, & Zhai, 2019). Hence, essential for the maintenance of subcellular NAD^+^ pools, NMNAT isoforms are localized to the different cellular compartments. In mammals, NMNAT1 is nuclear and NMNAT2 is cytoplasm-localized (Berger et al., 2005). In *Drosophila*, the Nmnat gene generates two protein isoforms through alternative splicing, nuclear Nmnat-PC and cytoplasmic Nmnat-PD (Ruan et al., 2015). Interestingly, we found in our Drosophila model, nuclear Nmnat is more active in promoting tumor growth; and nuclear NMNAT1 and cytoplasmic NMNAT2 have similar strong phenotypes in human glioma cells. Moreover, overexpressing nuclear NMNAT1 is slightly more efficient than cytoplasmic NMNAT2 to promote cell growth and proliferation. This finding has several implications. First, this suggests the requirement for nuclear NAD^+^-consuming events in tumor growth and the importance of supplying the nuclear pool of NAD^+^ on-demand by nuclear-localized NMNAT. Indeed, as our results show, NAD^+^-dependent PARylation and deacetylation of p53 underlies the mechanism of tumorigenesis. Second, the difference in the tumor-promoting effects of nuclear vs. cytoplasmic NMNAT isoforms may inform cellular metabolic needs and genotoxic load. Interestingly, the databases show NMNAT1 and NMNAT2 genes appear to be amplified in distinct cancer types (Figure 10-figure supplement 1 and 2). Future work is required to identify the specific roles of NMNAT1 and NMNAT2 in different cancer types. It is important to note that in mammalian cells, nuclear and cytoplasmic NMNATs can regulate each other’s activity, likely through feedback from dynamic pool of substrates NMN and ATP, as overexpressing cytoplasmic NMNAT may exhaust the supply of NMN therefore repress nuclear NAD^+^ synthesis (Ryu et al., 2018). Consequently, altering nuclear NAD^+^ pool may regulate gene transcription and influence cell differentiation or proliferation state (Ryu et al., 2018). Our observation of the specific upregulation of endogenous nuclear NMNAT upon oncogenic RAS-expression further supports the hypothesis that nuclear and cytoplasmic NMNAT react differently in stress conditions and likely be important in different stages of tumor growth.

### NAD^+^-mediated post-translational modifications of p53: a balancing act

PARylation, phosphorylation, acetylation, and ubiquitination are post-translational modifications that have been shown to regulate the stability and activity of p53 (Bode & Dong, 2004). Among the most common post-translational modifications of p53, PARylation and acetylation are both NAD^+^-consuming processes mediated by NAD+-dependent enzymes, PARPs, and SIRTs (Lee, Na, Kim, Lee, & Lee, 2012; Vaziri et al., 2001). When PARP1 activity is induced in the DNA damage response process, extensive protein PARylation occurs and many proteins including p53 and DNA repair machinery components are PARylated (Ame et al., 2004). Numerous studies have shown that PARylation of p53 may inhibit p53-mediated function including cell cycle arrest and apoptosis (Kanai et al., 2007; Simbulan-Rosenthal et al., 1999; Simbulan-Rosenthal et al., 2001). With abundant NAD^+^ supply, PARylation is an efficient way to repair DNA damage and ensure cell survival; while under conditions of insufficient NAD^+^ supply, apoptosis is induced (Herceg & Wang, 2001). In response to DNA damage, the activity of p53 is also modulated by acetylation. Acetyl-p53 is resistant to degradation by ubiquitination and has higher stability, and therefore can exert longer effects of growth arrest, senescence, and apoptosis (M. Li, Luo, Brooks, & Gu, 2002). NAD^+^-dependent PARylation and acetylation have the opposite effects on p53 activity, where PARylation inhibits p53 activity and acetylation prolongs p53 activity. NAD^+^ thus plays a critical role in balancing the pro-apoptotic activity of p53. NMNAT1 regulates functions of NAD^+^-dependent enzymes such as SIRT1 and PARP1 (Zhang et al., 2009; Zhang et al., 2012). Interestingly, our results identified a trimeric complex of p53-NMNAT-PARP1 and increase of PARP1 and SIRT1, which supports the model that NMNAT recruits NAD^+^-utilizing enzymes, including PARP1 and SIRT1 with protein substrates, and locally supply NAD^+^ for NAD^+^-dependent protein modification. Such a protein complex will not only sustain the local supply of NAD^+^ but also facilitate and expedite modification process.

It is important to note that p53 is not the only target for PARylation and deacetylation regulation. The role of NMNAT in PARylation of other target proteins has also been indicated. For example, it has been shown that decreased NMNAT1 expression caused nuclear NAD^+^ deficiency and subsequently reduced PARylation of multifunctional nuclear protein CCCTC-binding factor, leading to epigenetic silencing of tumor suppressor genes (Henderson et al., 2017). This report together with our findings support a specific role of nuclear NAD^+^ in modulating tumorigenesis through regulating posttranslational modifications including PARylation and deacetylation. Our findings in both *in vivo* and *in vitro* models highlight NMNAT’s roles in promoting glioma development. Specifically, the direct interaction we identified among p53, NMNAT, and PARP1 would have important implications in considering NMNAT as a potential target for glioma therapy. Because the protein-protein interaction interface of NMNAT/p53/PARP1 could provide allosteric targeting of NMNAT, in addition to its enzyme pocket, this may open new possibilities for alternative inhibitors of NAD^+^-dependent pathway with less toxicity.

In conclusion, our finding identified NMNAT as an NAD^+^ synthase that plays an essential role in regulating function and activation of p53 during DNA damage-induced apoptosis in cells. These results supported the development of specific NMNAT inhibitors as a potentially useful option for cancers with upregulated NMNAT levels.

## ACKNOWLEDGMENTS

We thank V. Chavez Perez and Q. Yang for technical expertise; and J Park for manuscript comments. We thank the FACS core facility at Sylvester Comprehensive Cancer Center, and Shannon Jacqueline Saigh for technical support. This research was supported by the Taishan Scholar Project of Shandong Province, the Science and Technology Support Program for Youth Innovation in Universities of Shangdong (2019KJM009), National Natural Science Foundation of China (82073888), Top Talents Program for One Case Discussion of Shangdong Province.

## AUTHOR CONTRIBUTIONS

Hongbo Wang, R. Grace Zhai, Jiaqi Liu, Xianzun Tao, and Yi Zhu, designed experiments; Zoraida Diaz-Perez, Chong Li and Kai Ruan generated critical reagents; Jiaqi Liu performed the experiments and prepared figures; R. Grace Zhai, Jiaqi Liu, Xianzun Tao, Yi Zhu, and Kai Ruan analyzed data. Jiaqi Liu, R. Grace Zhai, Chong Li, Xianzun Tao, Yi Zhu, and Kai Ruan wrote and edited the manuscript.

## MATERIALS AND METHODS

### Fly stocks and culture

Flies were maintained at 25 °C room temperature with standard medium. The following lines were used in this study obtained from the Bloomington *Drosophila* Stock Center: (1) The driver used in all experiments: *repo-GAL4*; (2) *UAS-Ras^v12^* (II); (3) *UAS-Ras^v12^* (III); (4) *UAS-Nmnat RNAi* (III). (5) *UAS-p35*; (6) *UAS-Diap1*; (7) *UAS-Dronc* RNAi; (8) *UAS-DCP1 RNAi*. UAS-*Drosophila melanogaster* Nmnat (*UAS-PC, UAS-PC^WR^, UAS-PD*) were generated in the laboratory.

### Fly treatment

Larvae were collected and treated with 100 μM of Pifithrin-α (Sigma-Aldrich, P4359) with standard medium at 25 °C room temperature.

### Human Glioma Cell Culture and treatment

T98G and U87MG (human glioma cells) cell lines were purchased from the American Type Culture Collection (ATCC, CRL-1609). SVG p12 cell line was from Dr. Michal Toborek (University of Miami). Cells were maintained in Eagle’s Minimum Essential Medium (EMEM, Sigma, M0325) supplemented with 10% Fetal Bovine Serum (FBS, ATCC, 30-2020). Cells were cultured at 37 °C, 5% CO_2_. To induce apoptosis, cells were treated with 50 μM of cisplatin for 8 hr (Sigma-Aldrich, 232120).

### Antibodies

The following commercially available antibodies were used: anti-Repo (1:250, DSHB, 8D12), anti-Caspase-3 (1:250 for Immunocytochemistry of fly brain, 1:1000 for Western blot analysis, Cell Signaling, 9665), anti-Cleaved Caspase-3 (1:1000, Santa Cruz, 9661), anti-H2AvD (1:50, Rockland, 600-401-914), anti-p53(E-5) (1:50, Santa Cruz, sc-74573), p53(DO-1) (1:1000, Santa Cruz, sc-126), anti-Drosophila Nmnat (1:3000), anti-NMNAT1 (1:1000, Abcam, ab45548), anti-NMNAT1 (1:1000, Santa Cruz, 271557), anti-NMNAT2 (1:500, Abcam, ab56980), anti-PARP1 (1:1000, Santa Cruz, sc-8007), anti-pADPr (1:1000, Santa Cruz, sc-56198), anti-SIRT1 (1:1000, Cell Signaling, 2492), anti-acetyl-p53 (1:1000, Cell Signaling, 2525), anti-β-actin (1:10,000, Sigma-Aldrich, A1978), anti-tubulin (1:300, Abcam, ab15246). The secondary antibodies conjugated to Alexa 488/546/647 (1:250, Invitrogen), or near-infrared (IR) dye 700/800 (1:5000, LI-COR Biosciences). HRP-anti-Mouse and HRP-anti-Rabbit (1:5000, Thermo fisher).

### Plasmid construction

Four recombinant plasmids were generated for this study: pDsRed, pDsRed-NMNAT1, pDsRed-NMNAT2.

### RNA interference

Small interference RNA sequences targeting human NMNAT were purchased (GenePharma). The siRNA sequences were listed in Supplementary (Fig 3-figure supplement 2).

### Real-time RT-PCR

The total RNA was extracted by TRIzol reagent (Invitrogen) from T98G cells according to the manufacturer’s protocol. cDNA was synthesized from RNA with a cDNA reverse transcription kit (Applied Biosystems). RNA was performed using a real-time system and SYBR green kit (Applied Biosystems). Relative gene expression was compared to actin as an internal control. The primers used in detection were listed in the Supplementary (Fig 3-figure supplement 2).

### Cells transfection

Cells for transfection were seeded in a 6-well culture vessel (VWR) containing EMEM media with 10% FBS for 24 hr. Plasmids or siRNA were transfected with transfection reagent (jetPRIME). Gene expression was measured by Western blot and Real-time qPCR after cells were transfected at 48 hr.

Cells were grown on 22-mm glass coverslips (VWR). After treatment, cells were rinsed three times with PBS, fixed for 15 min in 4% paraformaldehyde, washed three times with PBS, and permeabilized with 0.4% Triton X-100 in PBS for 5 min. After three times washing in PBS, blocking was performed by incubation in 5% normal goat serum in PBTX (PBS with 0.1% Triton X-100) at 37 °C for 30 min. Incubation with primary antibodies was performed in 5% goat serum in PBTX at 37 °C for 2 hr. Next, cells were washed three times with PBS and incubated for 1 hr at 37 °C with secondary antibodies in 5% goat serum in PBTX. Then, after three times washing with PBS, cells were stained with 4’,6diamidino-2-phenylindole (DAPI, 1:300, Invitrogen) at 37 °C for 5 min in PBTX solution. The cells were washed three times with PBS, and the coverslips were mounted on glass slides with VECTASHIELD Antifade Mounting Medium (Vector Laboratories) and kept at 4 °C before imaging.

### Immunocytochemistry of fly brain

The larval brains were dissected in phosphate-buffered saline (PBS, pH 7.4), and fixed in PBS with 4% formaldehyde for 15 min. After the brains were washed in PBS containing 0.4% (v/v) Triton X-100 (PBTX) for 15 min three times, the brains were incubated with primary antibodies diluted in 0.4% PBTX with 5% normal goat serum overnight. Then, secondary antibodies at room temperature for 1 h, followed by 4’,6diamidino-2-phenylindole (DAPI, 1:300, Invitrogen) staining for 10 min. Brains were mounted on glass slides with VECTASHIELD Antifade Mounting Medium (Vector Laboratories) and kept at 4 °C before imaging.

### Confocal image acquisition and image analysis

Confocal microscopy was performed with an Olympus IX81 confocal microscope coupled with x10, x20 air lens or x40, x60 oil immersion objectives, and images were processed using FluoView 10-ASW (Olympus). Specifically, Fig. 7B, C were analyzed using the Image J interactive 3D surface Plot plugin.

### Western blot analysis

Proteins were extracted from cells in either radioimmunoprecipitation assay (RIPA) buffer 1 mM protease inhibitor cocktail (Sigma-Aldrich). Samples were heated at 100 °C for 10 min in a 4X loading buffer. Proteins were separated on a Bis-Tris gel and transferred to nitrocellulose membranes. Then, membranes were blocked with blocking buffer (Rockland) for 1 hr at room temperature. Primary antibodies were incubated at 4 °C overnight and secondary antibodies were incubated for 1 hr at room temperature. Images were processed on an Odyssey Infrared Imaging System and analyzed using Image Studio software.

### Dot blot analysis

Proteins were extracted from cells in either radioimmunoprecipitation assay (RIPA) buffer 1 mM protease inhibitor cocktail (Sigma-Aldrich). Proteins were loaded on PVDF membranes. Then, membranes were blocked with Casine buffer for 1 hr at room temperature. Primary antibodies were incubated at 4 °C overnight and secondary antibodies were incubated for 1 hr at room temperature. Images were processed on an Amersham Imager 600 and analyzed using image J software.

### Cell proliferation test

Cells were seeded into the E-Plate 96 (ACEA) with the same confluence per well. Then, the Plate was incubated at 37 °C in 5% CO_2_ for about 100 hr. The instrument was used to monitor the cell growth index. The cell growth curve was drawn with the value of each group from xCELLigence RTCA SP instrument.

### Colony formation assay

Cells were seeded with 1000 per well in a 6-well plate containing 2 ml medium and replaced medium every two days. Cells were washed with 1 ml PBS three times and fixed with 1 ml formaldehyde for 15 min. After washed with PBS, cells were stained in 0.1% crystal violet buffer (Sigma) for 15 min. Cells were washed with pure water gently, and plates were put at room temperature to dry. Images were processed on an Amersham Imager 600 and analyzed using image J software.

### Immunoprecipitation

Proteins were extracted from cells in RIPA buffer. Proteins were incubated with Protein-A beads (Thermo Fisher Scientific) conjugated with anti-p53 antibody or Mouse IgG at 4 °C overnight with gentle shaking. After removed the supernatant, the bead pellets were collected and suspended with lysis buffer. Proteins were heated with loading buffer for 10 min at 100 °C for loading to gel.

### Flow Cytometry

Cells were prepared according to cell cycle and cell apoptosis detection kits (BD Pharmingen) after knockdown of NMNAT 72 hr.

### Statistics

For each statistical test, biological sample size (n), and P value are indicated in the corresponding figure legends. All data in this manuscript are shown as mean ± SD or median ± quartiles (specified in figure legends). t-test was used to compare between two groups, and one-way ANOVA with Bonferroni’s post hoc test was applied to compare among three or more groups. Data were analyzed with Prism (GraphPad Software). Specifically, fly survival data were analyzed by the Chi-square test in R.

## Supplemental Information

**Figure 1-figure supplement 1.**
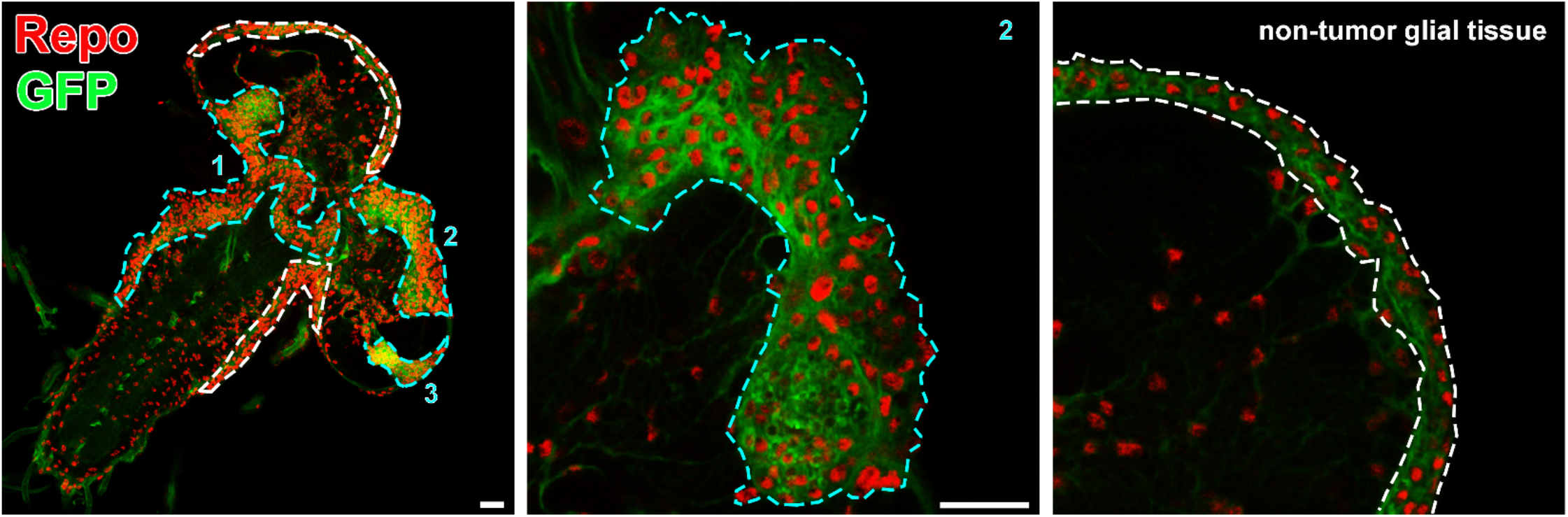
Glial neoplasia tissue area in *Drosophila* larval CNS. The *Drosophila* larval CNS with glial expression of Ras^v12^ + GFP (green) was probed for Repo (red). The glial neoplasia tissue and non-tumor glial area are marked with cyan and white dashed lines, respectively. Scale bars, 30 µm.

**Figure 2-figure supplement 1.**
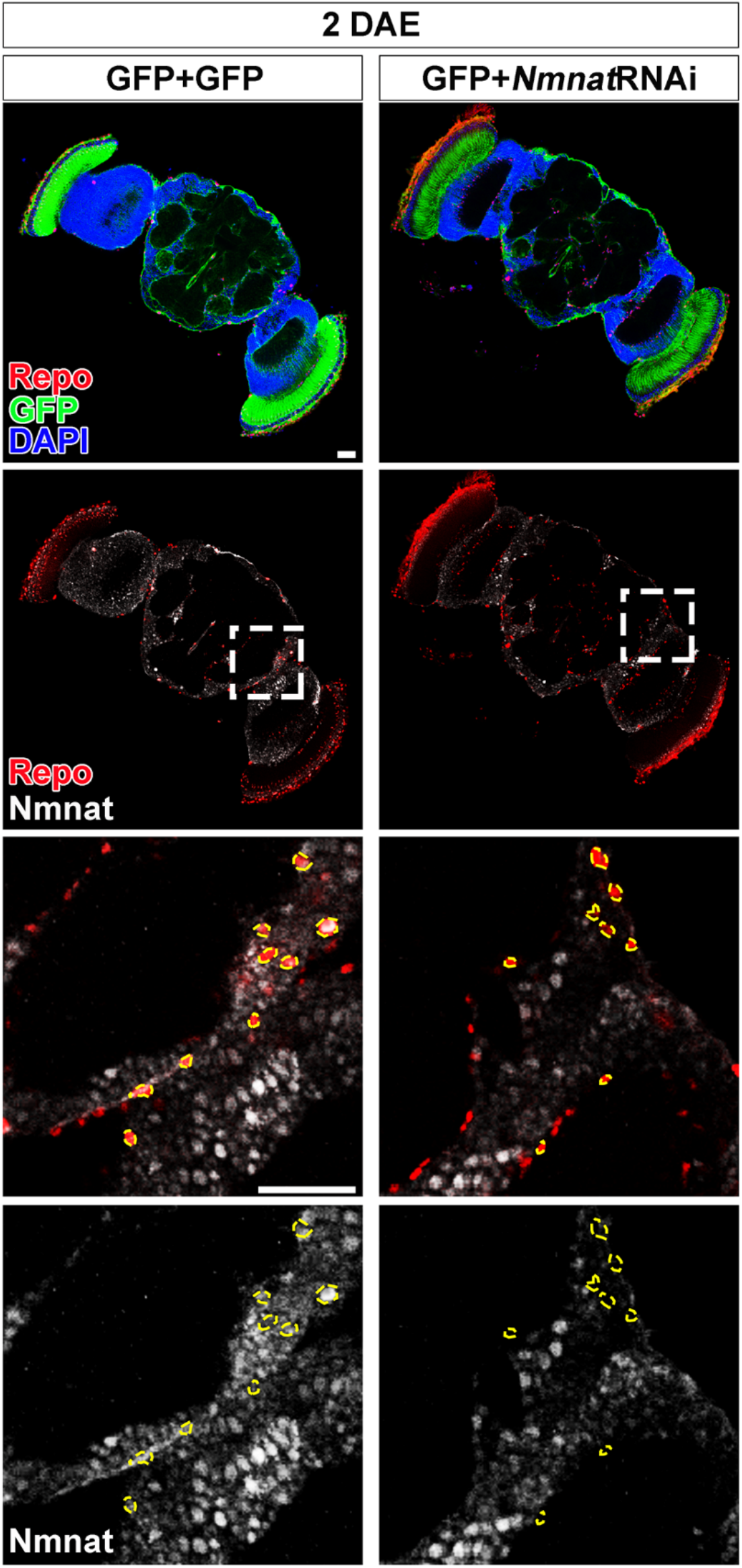
NMNAT downregulation in glial cells does not affect brain morphology in adult fly. Brains of 2 DAE flies with glial expression of GFP + GFP or GFP + NmnatRNAi were probed for Repo (red), Nmnat (white), and DAPI (blue). The third and fourth rows are high magnification of the boxed areas in the second row. Scale bars, 30 µm.

**Figure 2-figure supplement 2.**
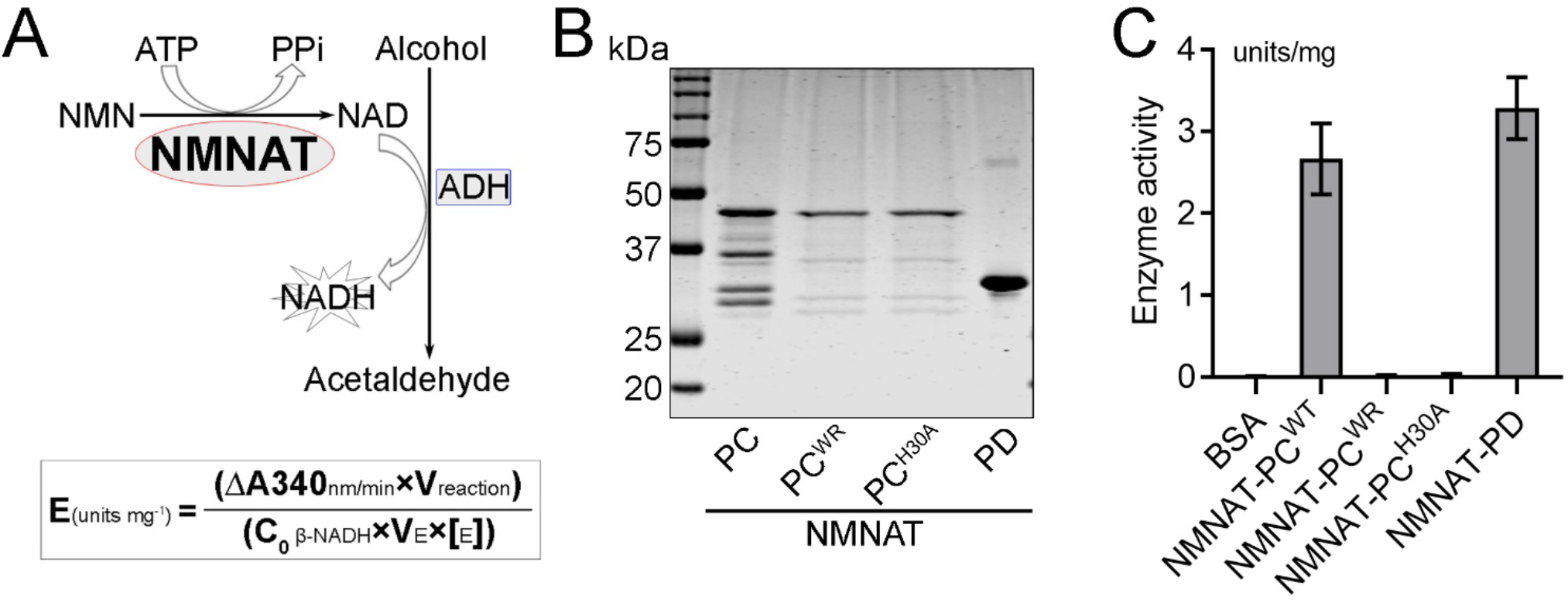
NMNAT-PC^WR^ has no NAD^+^ synthesis enzyme activity. (**A**) Diagram of the continuous coupled enzyme assay where NAD^+^ synthesized by NMNAT is reduced to NADH by alcohol dehydrogenase (ADH). The production of NADH is measured by absorbance at 340 nm. NMNAT activity (units per milligram of recombinant protein) is calculated from the linear progression curve by the formula at the bottom, where Coβ-NADH, the extinction coefficient of β-NADH at 340 nm, is 6.22. (**B, C**) NAD^+^ synthesis activity of recombinant NMNAT-PC^WT^, NMNAT-PC^WR^, NMNAT-PC^HA^, and NMNAT-PD (**B**) was measured by the continuous coupling assay as shown in A. Bovine serum albumin (BSA) was used as a negative control. Data are presented as mean ± s.d., n=4.

**Figure 2-figure supplement 3.**
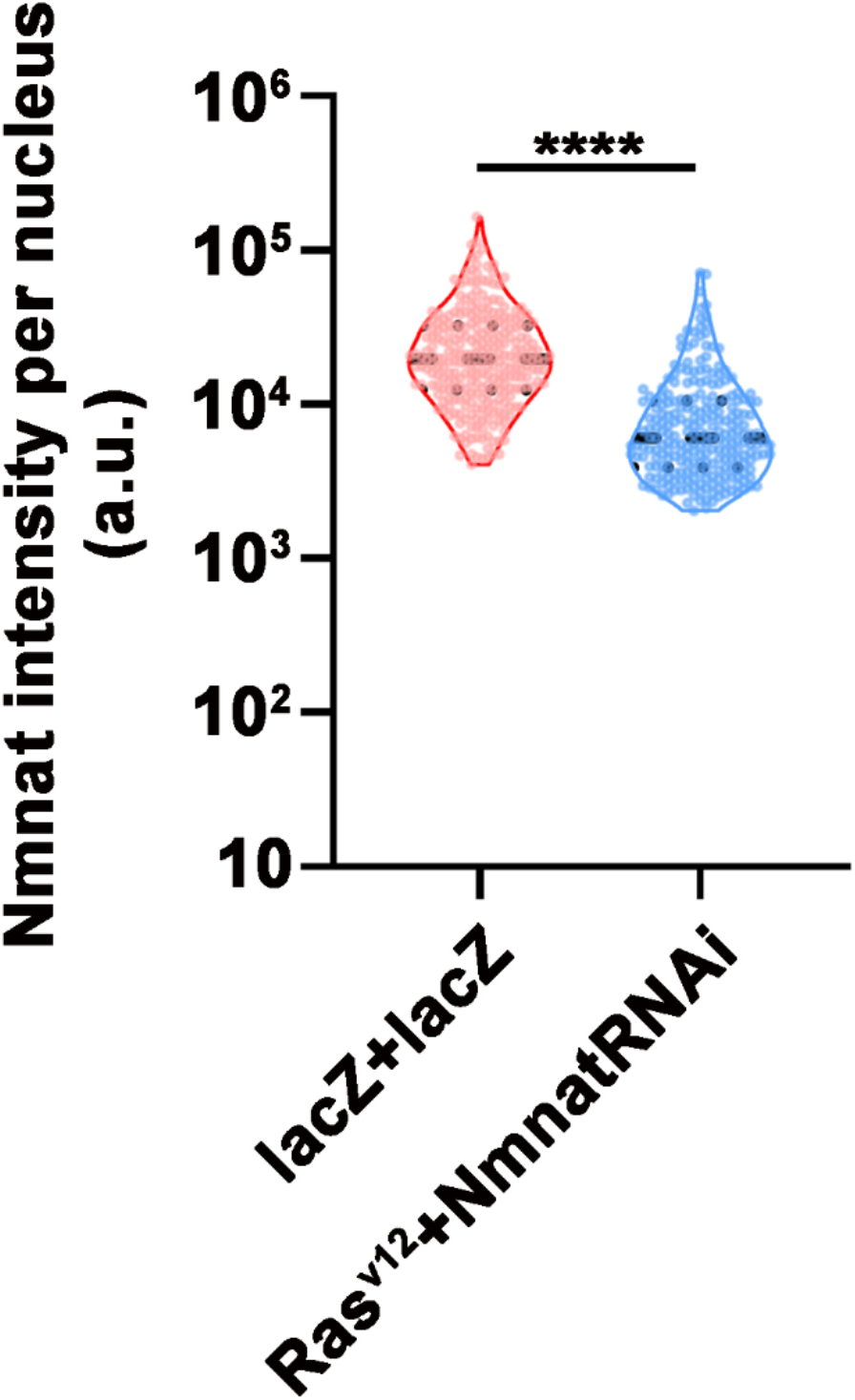
NMNAT expression is lower in NmnatRNAi fly than wild type control fly. Quantification of Nmnat intensity in glial cells or glial neoplasia area. Flies were expressing lacZ+lacZ or Ras^v12^+NmnatRNAi. Data are presented as median ± quartiles, n ≥ 3. Significance level was established by one-way ANOVA post hoc Bonferroni test. \*\*\*\**P* ≤ 0.0001.

**Figure 3- figure supplement 1.**
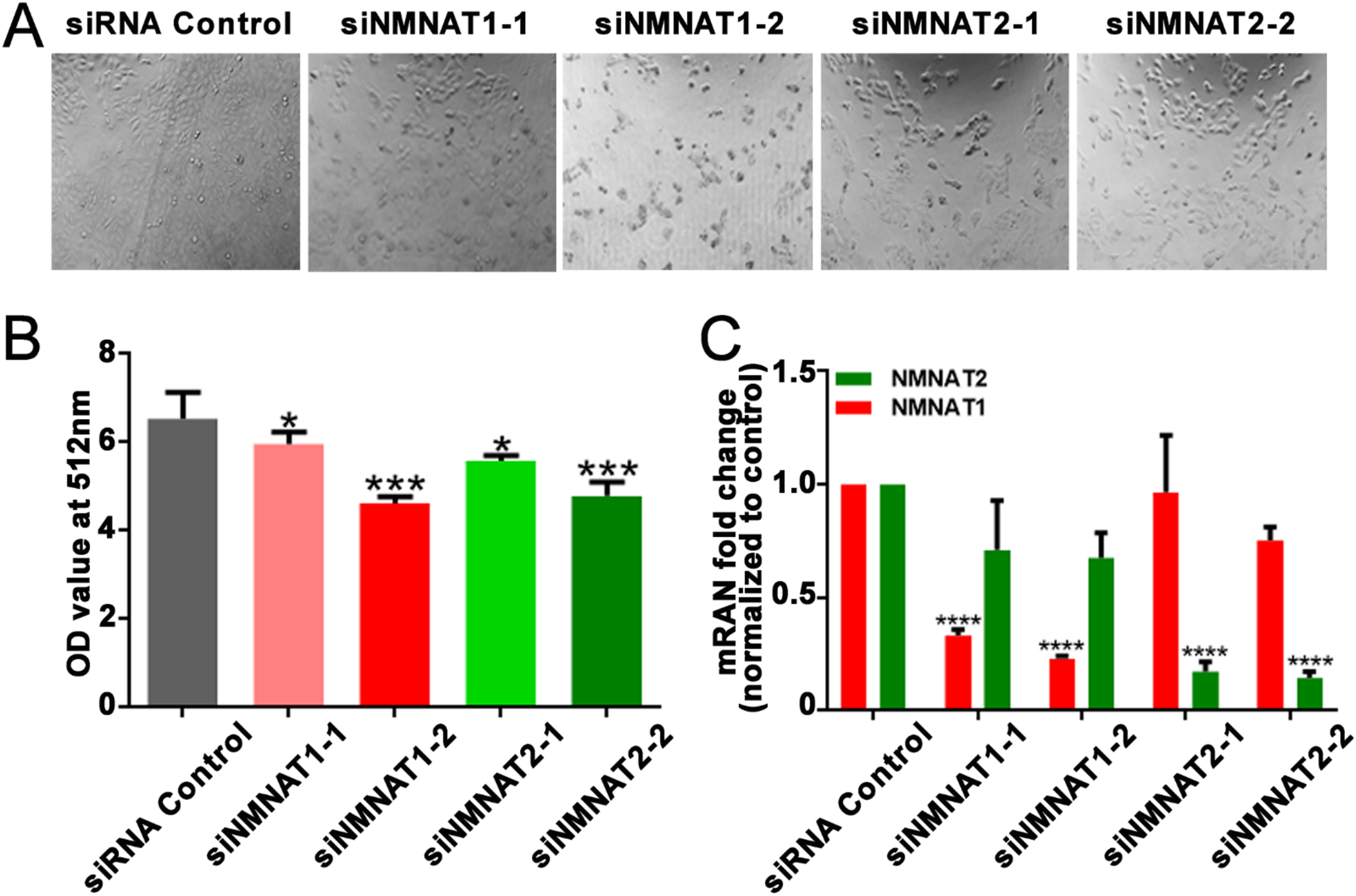
T98G cells viability is inhibited after knockdown NMNAT1 or NMNAT2. (**A**) The confluency of T98G cells is decreased after transfected with siNMNAT1 or siNMNAT2 96 hours. (**B**) Cell viability 96 hours after transfection was measured by an MTT assay. Data are presented as mean ± s.d., n ≥ 3. Significance level was established by one-way ANOVA post hoc Bonferroni test. (**C**) NMNAT1 and NMNAT2 transcript levels after siRNA transfection 72 hours. Data are normalized to siRNA control group. Data are presented as mean ± s.d., n ≥ 3. Significance level was established by one-way ANOVA post hoc Bonferroni test. \**P* ≤ 0.05. \*\*\**P* ≤ 0.001. \*\*\*\**P* ≤ 0.0001.

**Figure 3- figure supplement 2.**
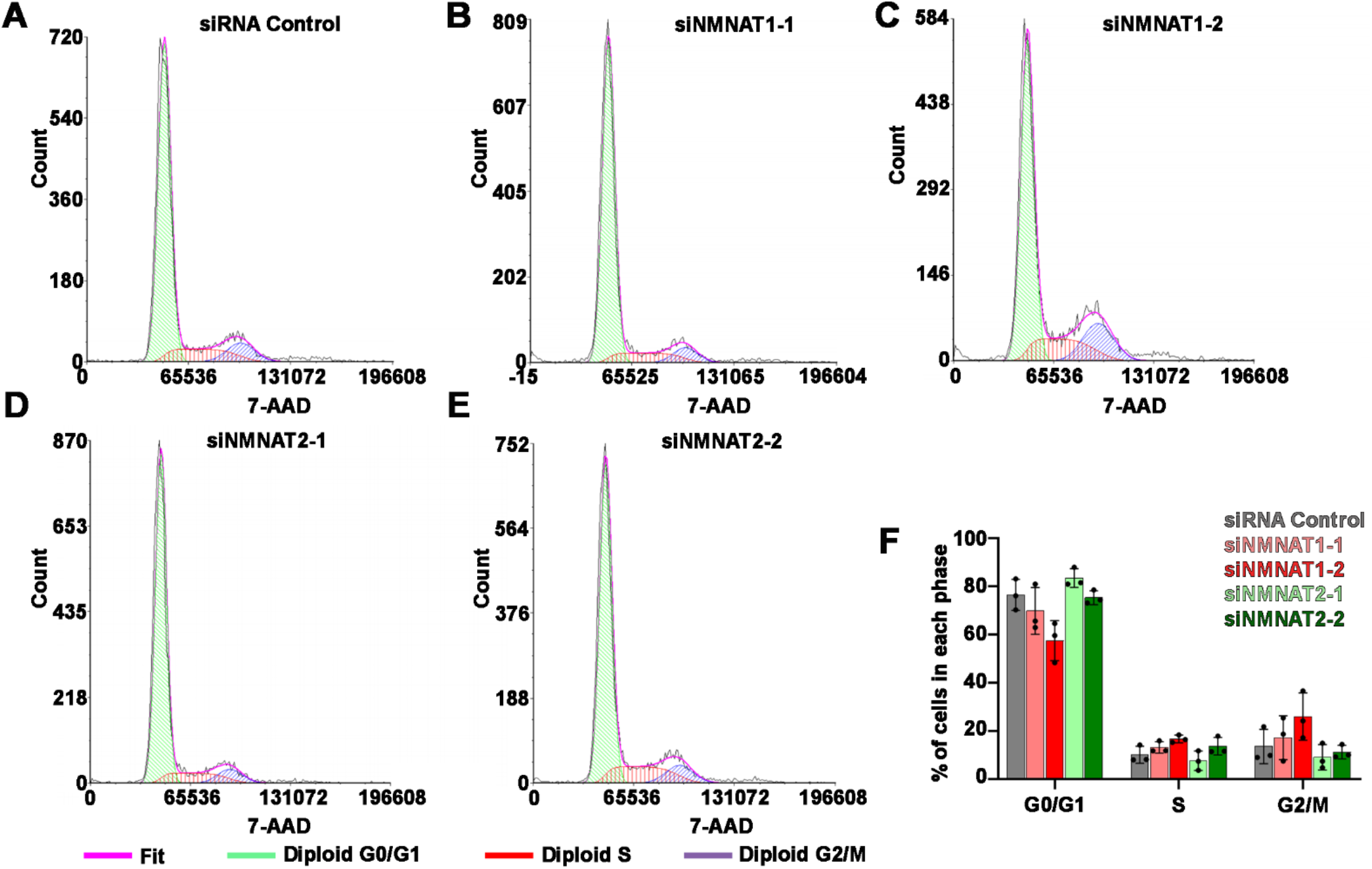
Knockdown of NMNAT does not affect cell cycle. **(A-E)** Cell cycle was detected by flow cytometry after T98G cells was transfected with siRNA Control, siNMNAT1- 1, siNMNAT1-2, siNMNAT2-1 and siNMNAT2-2 respectively. **(F)** Quantification of cells in each cell cycle phase. Data are presented as mean ± s.d., n = 3. Significance level was established by one-way ANOVA post hoc Bonferroni test.

**Figure 3- figure supplement 3.**
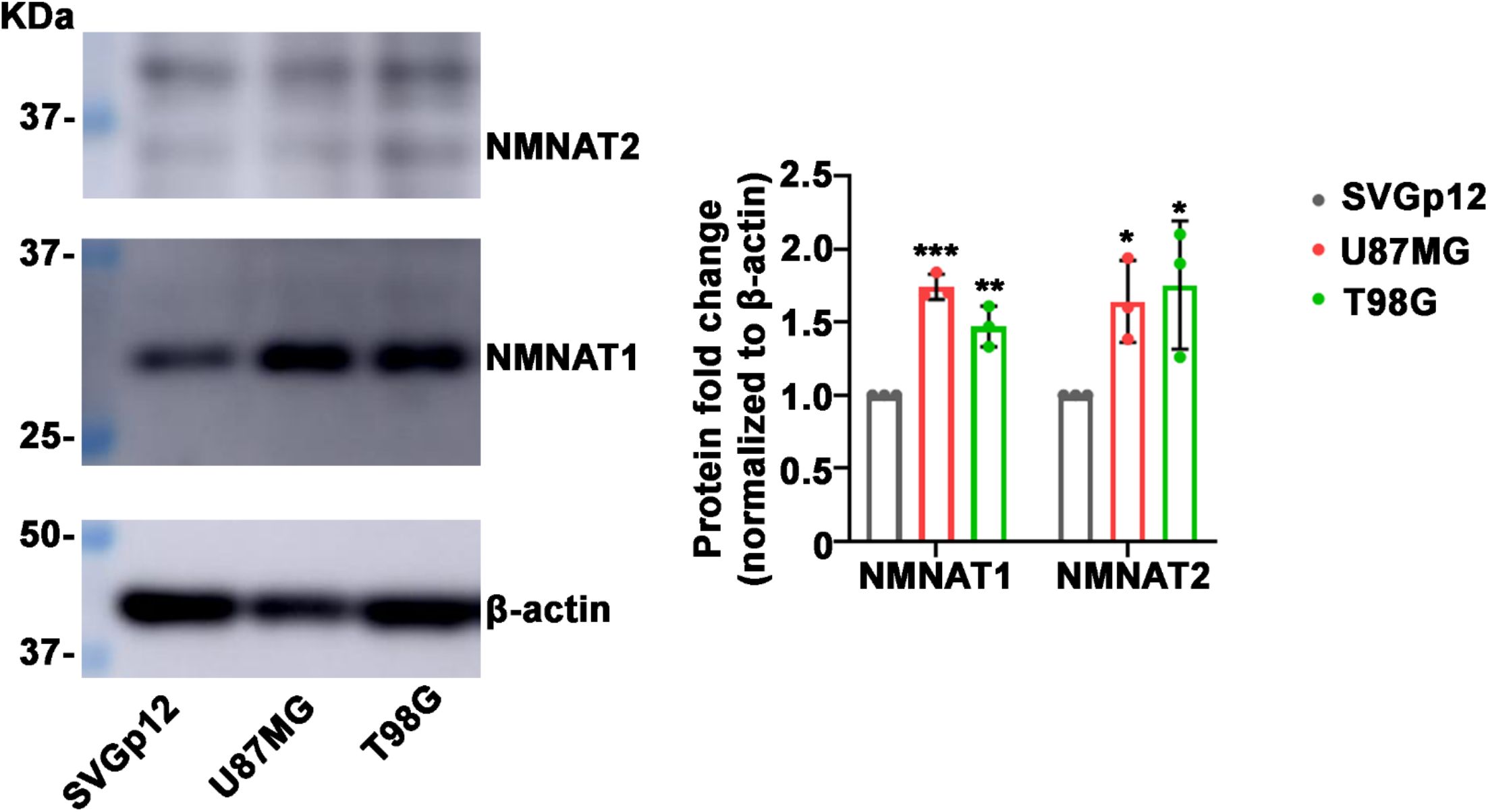
NMNAT protein is upregulated in human glioma cells. SVG p12 is human glial cells. T98G and U87MG are human glioma cells. Proteins were extracted from cells were probed for NMNAT1, NMNAT2 and β-actin and quantification. Data are presented as mean ± s.d., n = 3. Significance level was established by t-test. \**P* ≤ 0.05. \*\**P* ≤ 0.01. \*\*\**P* ≤ 0.001.

**Figure 3-table supplement 1.**
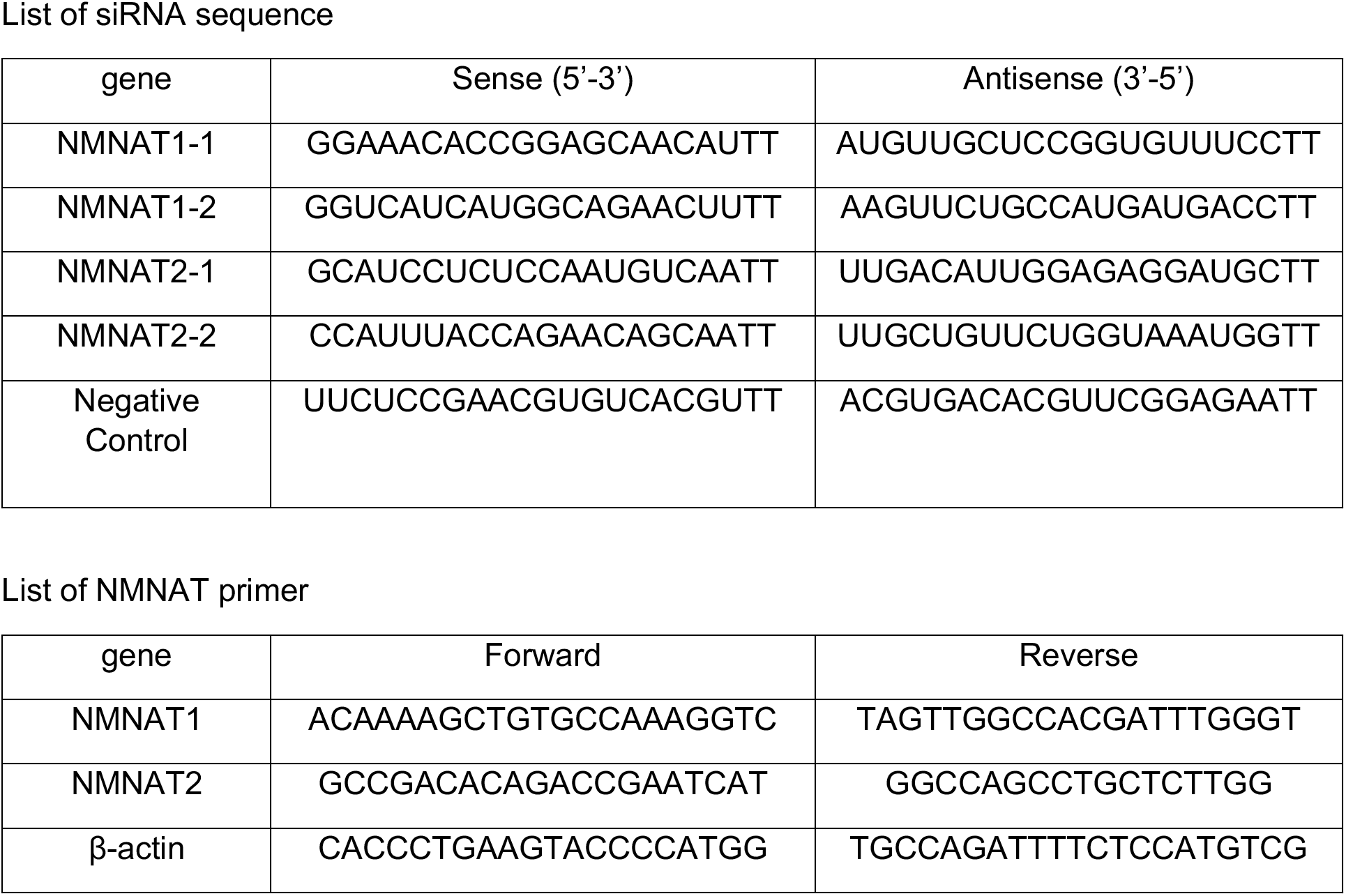
siRNA sequences for NMNAT1/2 knockdown and primer sequences for PCR.

**Figure 4- figure supplement 1.**
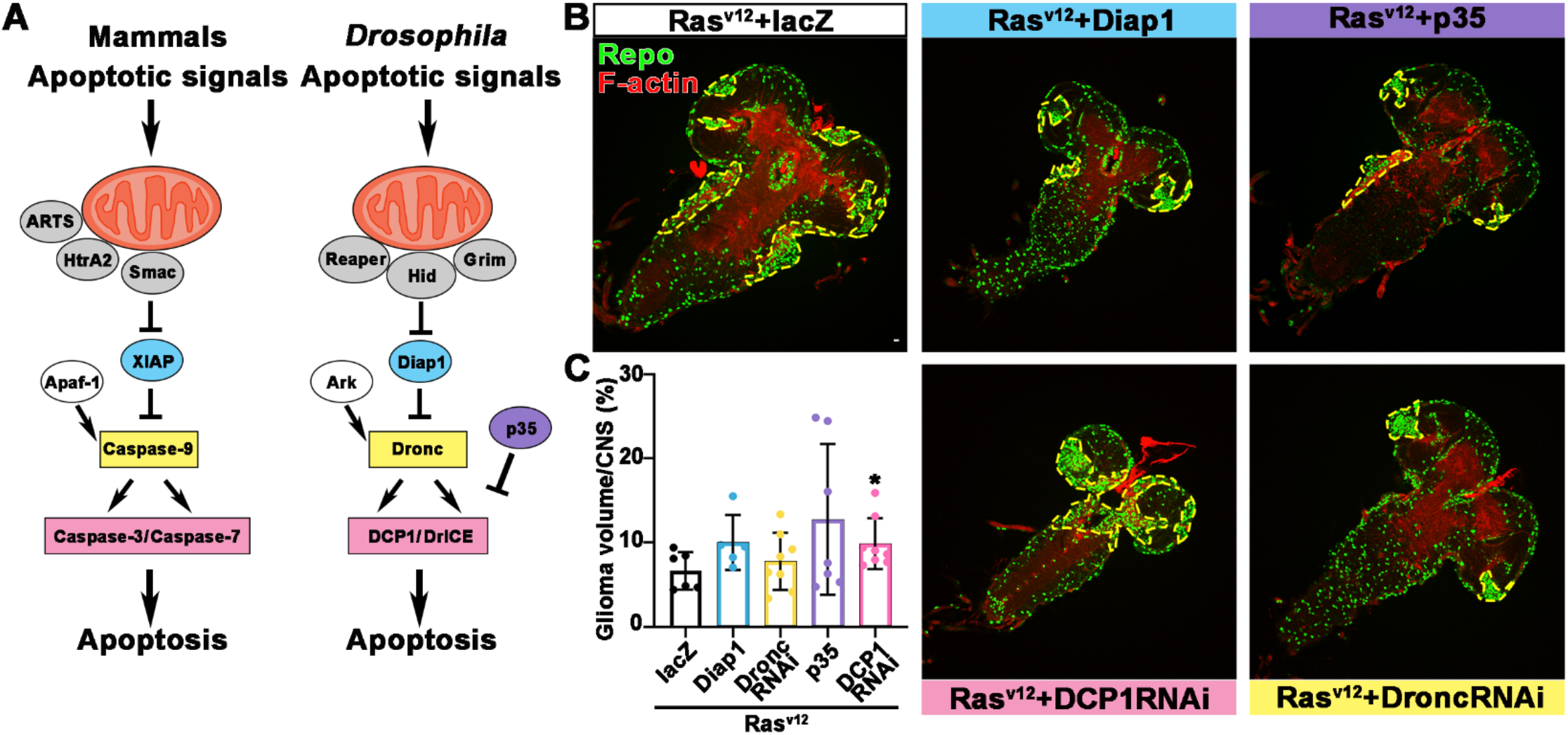
Blocking caspase pathway in *Ras^v12^* overexpressing fly. **(A)** Diagram of caspase pathway in mammalian and *Drosophila*. **(B)** Flies with Ras^v12^+lacZ, Ras^v12^+Diap1, Ras^v12^+p35, Ras^v12^+DCP1RNAi and Ras^v1^2+DroncRNAi were probed for Repo (green) and F-actin (red). (**C**) Quantification of ratio of glial neoplasia volume in CNS. Data are presented as mean ± s.d., n > 3. Significance level was established by t-test. \**P* ≤ 0.05.

**Figure 5-figure supplement 1.**
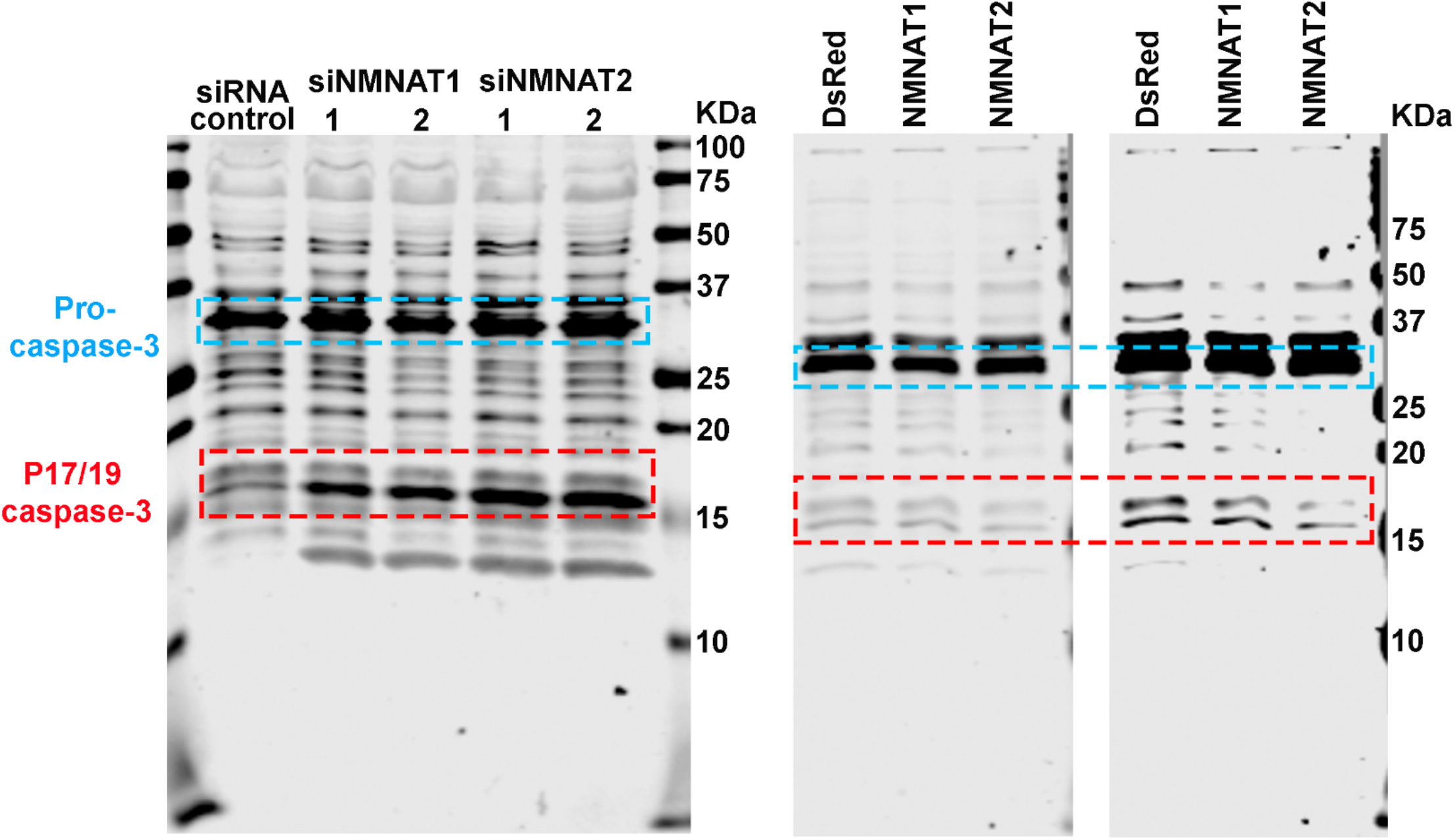
Membrane of Western blot. Proteins were extracted from T98G cells transfected with siRNA (left), or plasmids and treated with cisplatin 8 hours (right) for western blot analysis. P32 was considered as pro- caspase-3. P17/19 was considered as cleaved caspase-3.

**Figure 5-figure supplement 2.**
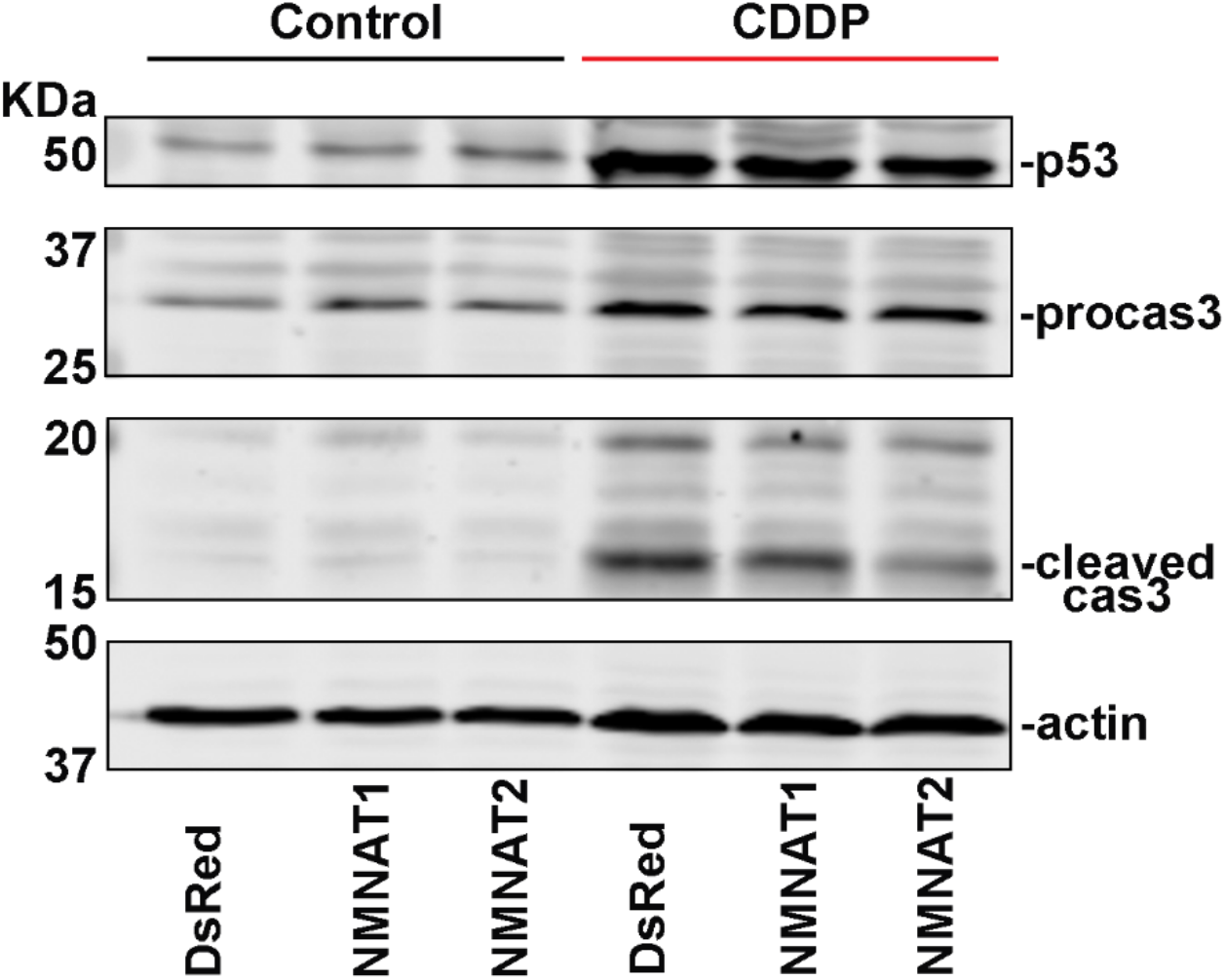
Cleaved Caspase-3 is reduced after NMNAT overexpression. U87MG cells were treated with CDDP and probed for p53, caspase-3 and β-actin.

**Figure 8-figure supplement 1.**
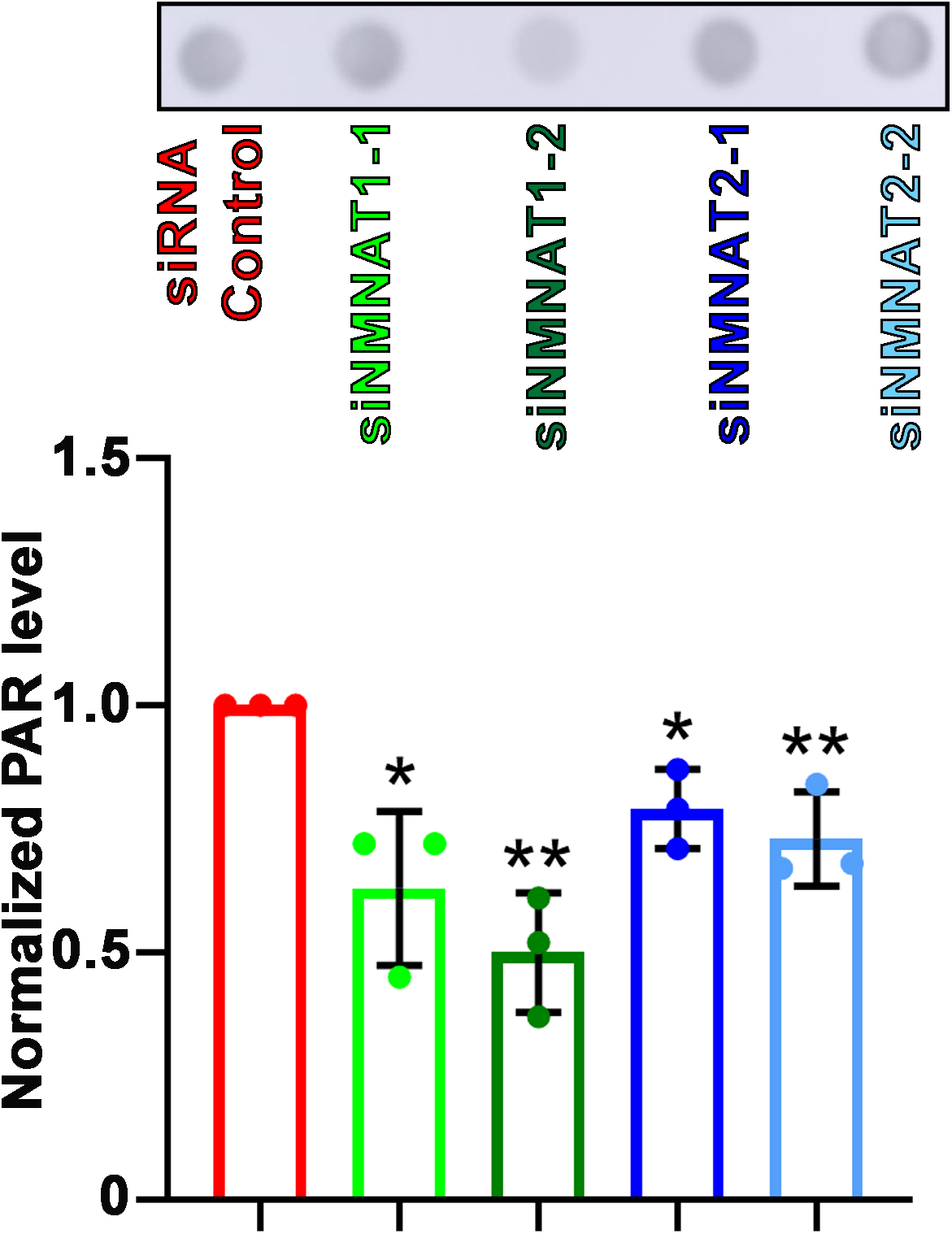
PARylation is reduced after NMNAT knockdown. Proteins were extracted from T98G cells transfected with siRNA for dot blot analysis using anti-PAR antibody and quantification normalized with β-actin as internal control. Data are presented as mean ± SD, n = 3. Significance level was established by t-test. \**P* ≤ 0.05. \*\**P* ≤ 0.01.

**Figure 9- figure supplement 1.**
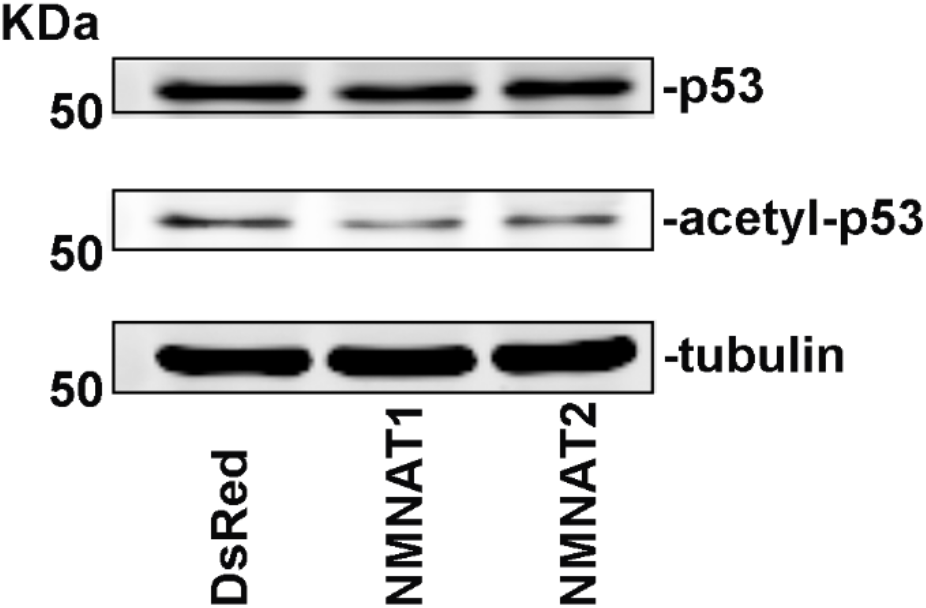
Acetyl-p53 is reduced after NMNAT overexpression in U87MG. U87MG cells were treated with CDDP and probed for p53, acetyl-p53 and tubulin.

**Figure 10- figure supplement 1.**
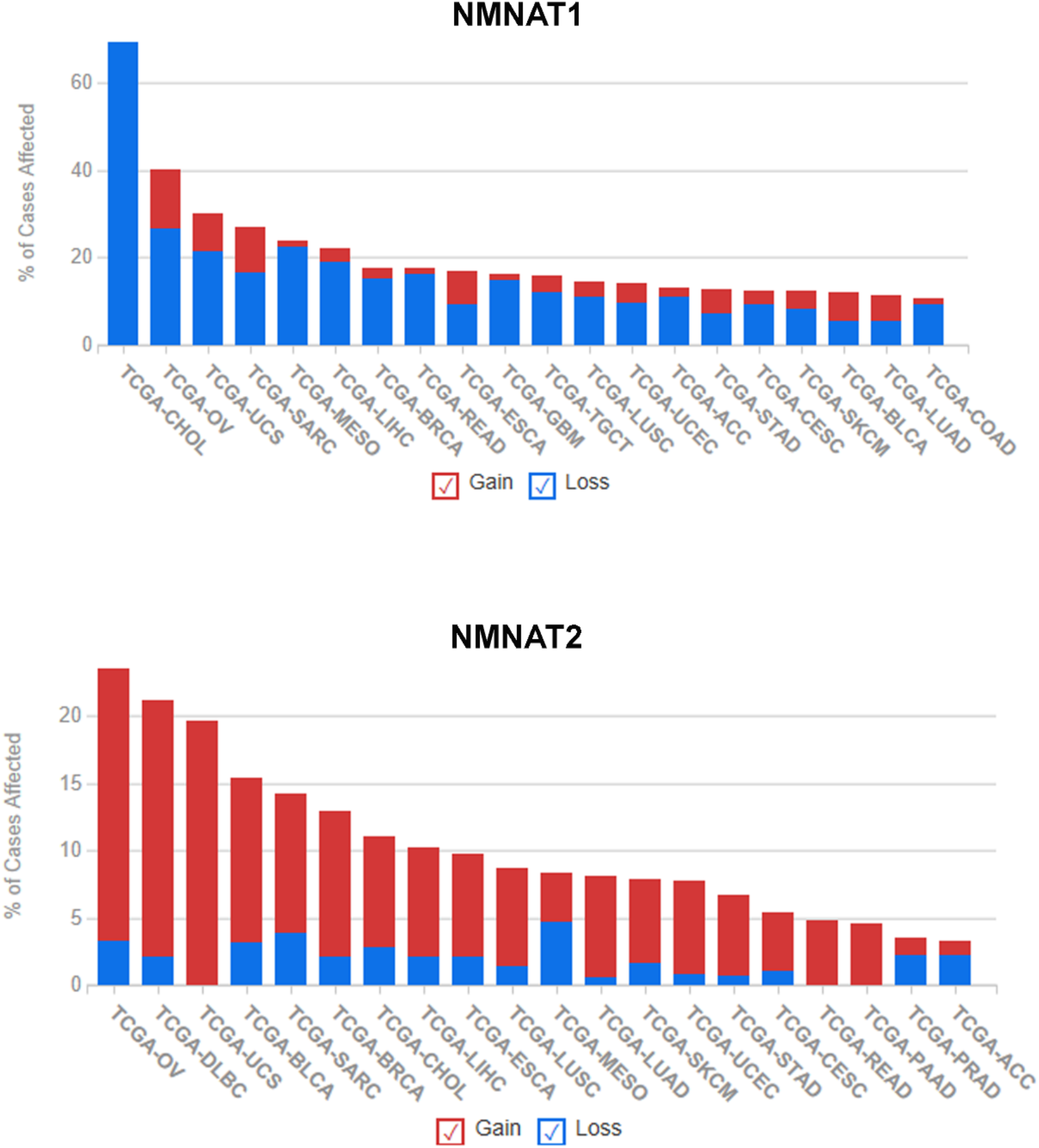
Summary of NMNAT1 and NMNAT2 alteration frequency in cancer types. Alteration of human NMNAT1 and NMNAT2 was queried in TCGA database (www.TCGA.com). 1522 cases with altered NMNAT1 and 931 cases with altered NMNAT2 across 32 projects are shown.

**Figure 10- figure supplement 2.**
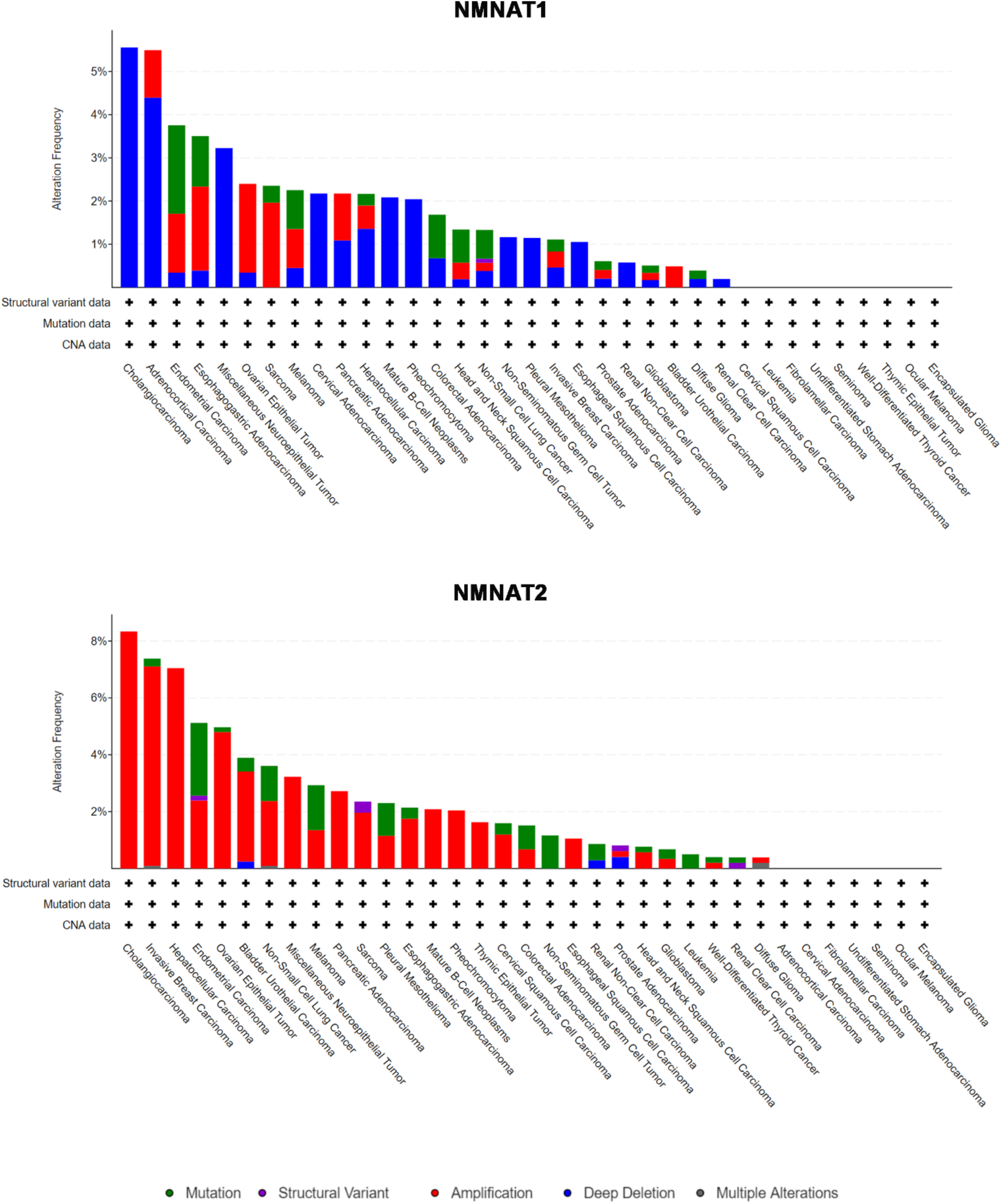
Summary of NMNAT1 and NMNAT2 alteration frequency in cancer types. Alteration of human NMNAT1 and NMNAT2 was queried in cBioPortal database (www.cbioportal.com) separately. 10967 samples in 35 cancer types are shown.

